# Studying a human genetic model of lung squamous cell carcinoma with organotypic cultures and xenografts uncovers distinct advantages of each system and implicates *NOTCH1* loss in tumour development

**DOI:** 10.1101/2025.08.28.672595

**Authors:** Julia Ogden, Robert Sellers, Anthony Oojageer, Sudhakar Sahoo, Caroline Dive, Carlos Lopez-Garcia

## Abstract

Selecting appropriate experimental systems is crucial in cancer research, where factors such as model relevance, cost, and resource availability guide decisions. A detailed understanding of the strengths and limitations of each model helps ensure their optimal use.

We recently developed a human lung squamous cell carcinoma (LUSC) model using genetically engineered human bronchial epithelial cells (hBECs). These were studied through organotypic air-liquid interface (ALI) cultures and standard in vitro assays, including proliferation, invasion, and anchorage-independent growth. However, we did not evaluate whether the same mutant hBECs behaved similarly in vivo, or if in vivo models offered distinct advantages.

To address this, we conducted a comparative phenotypic analysis of mutant hBECs derived from the same donor in both ALI cultures and xenografts in immunocompromised mice. Both models followed a similar oncogenic trajectory, involving squamous differentiation and activation of Nrf2 and PI3K/Akt pathways, characteristic of the classical LUSC subtype.

However, some transcriptomic differences related to an increase in microtubule formation and cell motility in xenografts emerged. Additionally, xenograft gene expression profiles more closely matched classical LUSC tumours than ALI cultures.

Importantly, we observed spontaneous squamous differentiation in the absence of *SOX2* overexpression and detected selection for *NOTCH1* mutations in specific in vivo mutants. Truncation of *NOTCH1* promoted squamous differentiation and suppressed mucociliary features in ALI cultures, underscoring its role as a potential LUSC driver.

In summary, mutant hBECs in vitro and in vivo showed largely consistent phenotypes, validating both systems. However, in vivo models can enable the unbiased discovery of new genetic LUSC driver genes. This highlights the complementary value of integrating both model types in LUSC research.

## INTRODUCTION

The development of tractable and robust cancer models relies on a transparent, detailed, and critical evaluation of their advantages and limitations for use in both basic and translational research. Specifically, recognizing the differences between experimental models and patient tumours is essential for their accurate application and for guiding model improvement. Additionally, factors such as scalability, versatility, and cost-effectiveness—without compromising biological relevance—must be considered when selecting experimental systems. Choosing the most informative models (e.g., mouse vs. human, in vitro vs. in vivo, 2D vs. 3D, artificial vs. patient-derived) requires balancing biological robustness with experimental feasibility.

We recently described a strategy for modelling lung squamous cell carcinoma (LUSC) through the genetic manipulation of human bronchial epithelial cells (hBECs) in vitro (1). This approach demonstrated that the activation of squamous differentiation, Nrf2, and PI3K/Akt pathways is required to transform hBECs. We also identified downstream processes validated in patient cohorts and proposed an evolutionary trajectory for LUSC development.

However, our previous work focused solely on in vitro models, and the behaviour of mutant hBECs in vivo was not explored. In vivo systems offer distinct advantages, including interactions with stromal components, long-term tumor growth, and selective pressures from the tumour microenvironment, which can reveal novel LUSC drivers. Conversely, some genotypes may not form tumours in vivo, limiting detailed analyses similar to our in vitro work.

Despite these considerations, a systematic comparison of in vivo and in vitro behaviour is essential for informed use of this model in cancer research. While other studies that reported genetic models of cancer by genetic manipulation of non-immortalised primary human cells (2–4) investigated tumourigenesis in vitro and in vivo but did not carry out a detailed comparison of both systems. Several studies have compared human cancer models in both contexts (5–7), but they focus on therapy responses in patient-derived models, not the identification of cancer drivers and developmental stages as we do. Moreover, no such comparative studies have been conducted in LUSC.

Therefore, we performed a side-by-side phenotypic analysis of mutant hBECs using the same genetic manipulations in organotypic air-liquid interface (ALI) cultures and xenografts in immunocompromised mice. Our findings showed similar results in vivo and in vitro regarding the requirements of activation of the three pathways for the transformation of hBECs and revealed transcriptomic differences that suggested microtubule formation and cell motility in vivo. Notably, we also identified an alternative LUSC development mechanism involving squamous differentiation driven by genetic inactivation of *NOTCH1*.

## RESULTS

### Macroscopic and histological analysis of mutant hBECs in vivo

To perform our comparative analyses, we generated the same nine hBEC genotypes used in our previous study (Figure 1A and Supplementary Figure 1A) (1), creating both ALI cultures and xenografts in immunocompromised NSG mice (Figure 1B). All models were derived from the same donor to ensure maximal reproducibility. These nine mutants harbour up to five genomic alterations (Figure 1A), targeting ubiquitous tumour suppressors commonly altered in LUSC (*TP53* and *CDKN2A* truncations) and components of the three most frequently dysregulated pathways in LUSC (8): the squamous differentiation, Nrf2, and PI3K/Akt pathways (*SOX2* overexpression and truncations in *KEAP1* and *PTEN*).

**Figure 1:**
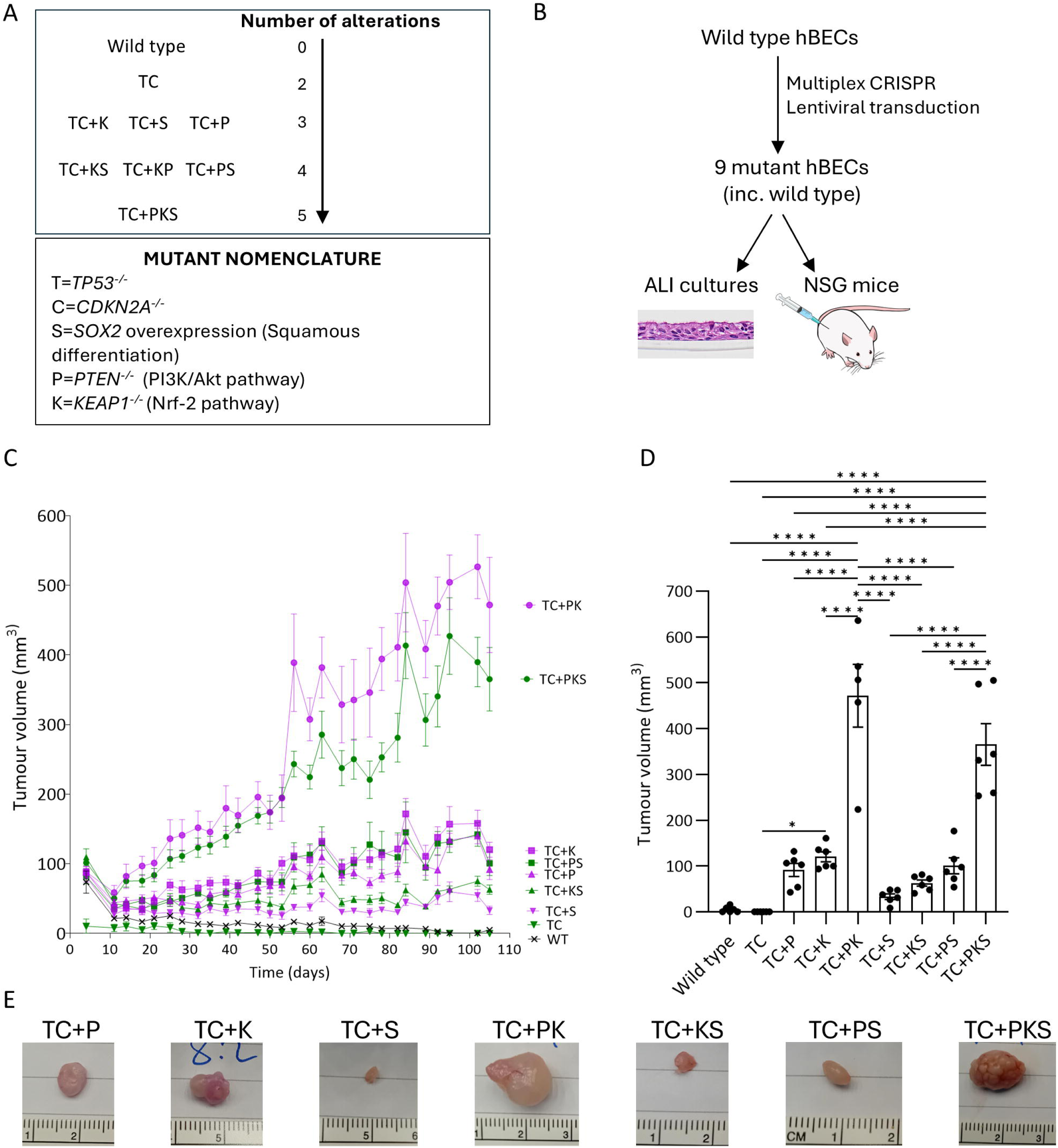
Experimental design and macroscopic features of xenografts generated by subcutaneous injection of genetically-engineered hBECs. **A.** Combinatorial strategy of genetic manipulation in hBECs designed to evaluate the individual and combined effect of the three most frequently dysregulated pathways in LUSC. **B.** Experimental design for a side-by-side comparison of the phenotypes of mutant hBECs in vitro (ALI cultures) and in vivo (subcutaneous injection in NSG mice). **C.** Xenograft growth curves for each hBEC genotype. Measurements of xenograft volume were taken every 3-4 days. Data is shown as mean±SEM (n=6 mice, except for TC+PK group, as one replicate of this group reached the maximum volume on day 89). **D.** Xenograft volumes of xenografts harvested on day 105. Data is shown as mean±SEM (n=5 mice for the TC+PK group, n=6 mice for the other groups). Adjusted p-values were calculated using one-way ANOVA followed by Holm-Šídák’s multiple comparisons test. Only significant comparisons (adjusted p-values<0.05) are shown. **E.** Representative macroscopic images of intact harvested xenografts. Statistical significance shown as: ∗p < 0.05, ∗∗p < 0.01, ∗∗∗p < 0.001, and ∗∗∗∗p < 0.0001

Preliminary experiments using intrathoracic injection failed to produce orthotopic tumour growth, prompting us to use subcutaneous transplantation. This method enables larger tumour volumes and facilitates the detection of somatic alterations relevant to LUSC development.

Subcutaneous nodules were allowed to grow for 105 days, except one TC+PK replicate, which was culled on day 89, as the nodule reached its maximum size. Analysis revealed that TC+PK and TC+PKS xenografts exhibited the fastest growth (Figure 1C) and reached the largest sizes (Figure 1D, E), while wild-type and TC mutant hBECs showed negligible growth (Figure 1C), with no visible nodules at the injection site. Although comparing xenograft size with ALI culture thickness is complex—due to insert size limits and the hollow structure of xenografts—we performed correlation analyses. Across all genotypes, there was no significant correlation between xenograft size and ALI thickness (Pearson r = 0.311, p = 0.4153; Supplementary Figure 1B). However, upon excluding the outlier TC+PK genotype, the correlation became significant (Pearson r = 0.7836, p = 0.0214; Supplementary Figure 1C), suggesting generally good concordance between in vivo and in vitro growth, except for TC+PK.

Histological comparisons between xenografts and ALI cultures revealed that TC+K and TC+PK xenografts displayed cystic structures with one or two lumens filled with mucinous fluid (Figure 2A, Supplementary Figure 2A), surrounded by a pseudostratified, bronchial-like epithelium positive for human mitochondria (Supplementary Figure 2B) and exhibiting mucociliary differentiation (Supplementary Figure 3A, B). These features were also present in their ALI culture counterparts (Figure 2B).

**Figure 2.**
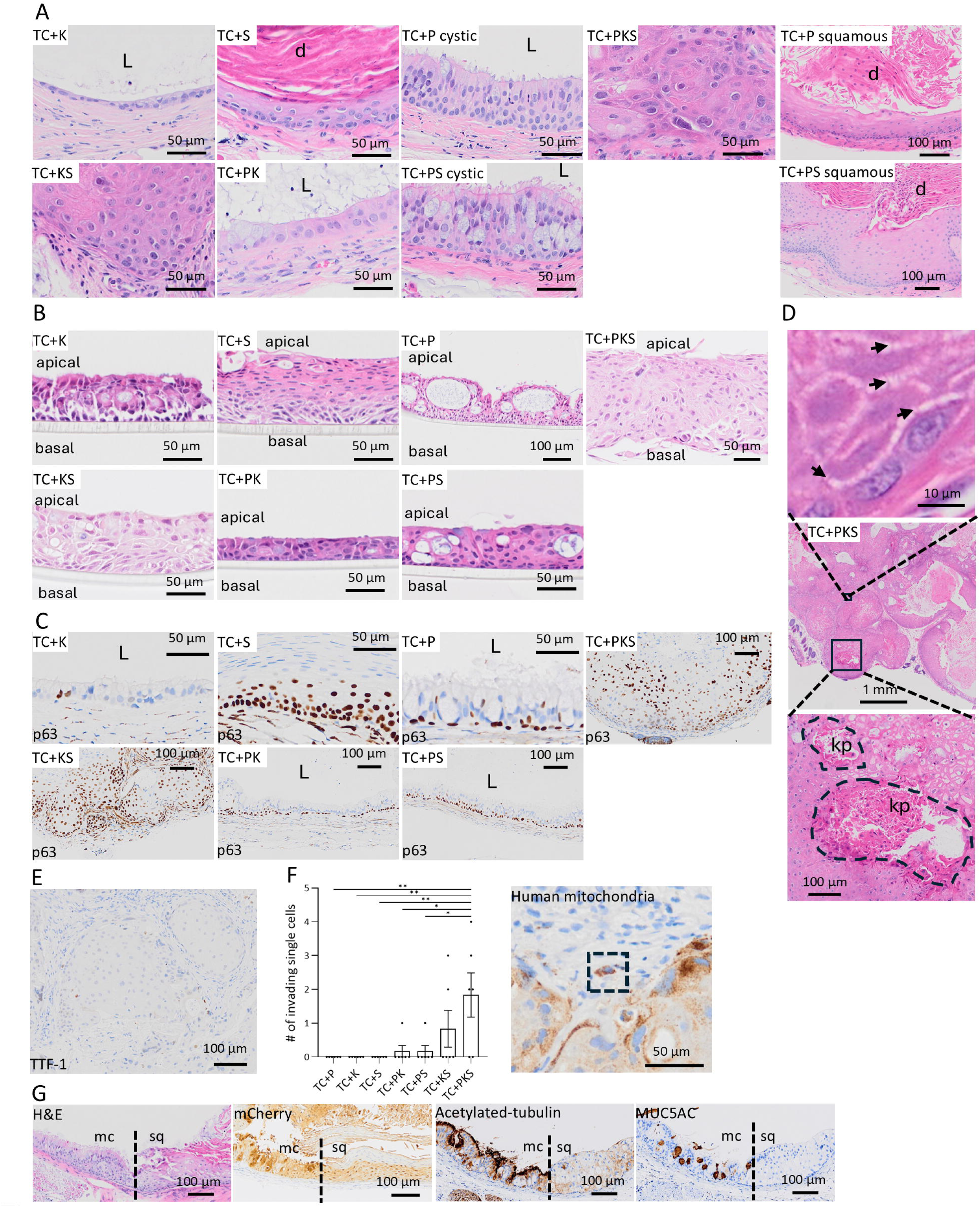
Comparative histological analysis of xenografts and ALI cultures generated with mutant hBECs from the same donor. **A.** Representative H&E images of the epithelial structure of xenografts. Images were taken from areas that comprise surrounding mouse stroma, epithelial layer and lumen for xenografts with cystic structure. For solid xenografts, images were taken from areas that comprise surrounding mouse stroma and tumour nests. For TC+P and TC+PS mutants, representative images of areas with squamous and mucociliary morphologies are shown separately. L=Lumen; d= dyskeratosis. **B.** Representative H&E images of the epithelial structure of ALI cultures. **C.** Representative images of p63 immunohistochemical staining of xenografts. L=Lumen. **D.** Images depicting the presence of intercellular bridges and keratin pearls in TC+PKS mutants, two features of well-differentiated LUSC. Arrows mark the presence of intercellular bridges. kp= keratin pearl. **E.** TTF-1 immunohistochemical staining of a TC+PKS xenograft showing the absence of expression of this lung adenocarcinoma marker. **F**. Quantification of total invading single cells into the mouse stroma and an example image of an invading single cell stained for human mitochondria. Data shown as mean±SEM (n=6 xenografts). Adjusted p-values were calculated using one-way ANOVA followed by Tukey’s multiple comparisons test (only significant comparisons are shown) **G.** Images depicting a xenograft area with adjacent mucociliary and squamous morphology in a TC+PS mutant. Images show H&E staining, and immunohistochemical staining for mCherry, acetylated-tubulin and MUC5AC. Areas with mucociliary differentiation show expression of acetylated-tubulin (cilia) and MUC5AC (goblet cells). Mc=mucociliary; sq=squamous. Statistical significance shown as: ∗p < 0.05, ∗∗p < 0.01

In contrast, TC+S, TC+KS, and TC+PKS xenografts showed predominantly solid architecture (Figure 2A, Supplementary Figure 2A). TC+S xenografts displayed polarized squamous stratified differentiation with marked apical dyskeratosis and absence of mucociliary differentiation (Figure 2A, Supplementary Figures 2A and 3A, B), mirroring their ALI counterparts (Figure 2B) and previous findings (1). TC+KS and TC+PKS xenografts exhibited disorganized solid growth with tumour nests positive for human mitochondria, surrounded by mouse stroma (Supplementary Figure 2B), and negative for mucociliary markers. These regions were positive for p63 (Figure 2C), contained keratin pearls and intercellular bridges—hallmarks of well-differentiated LUSC (Figure 2D)—and were negative for the lung adenocarcinoma marker TTF-1 (Figure 2E).

Another key finding of our in vitro work was the acquisition of an invasive phenotype that was most prominent in the TC+PKS mutant. In line with those results in vitro, TC+PKS xenografts showed the highest frequency of invasive cell into the mouse stroma (Figure 2F).

TC+P xenografts showed greater histological divergence from their ALI counterparts. While TC+P ALI cultures presented with hyperplasia and intraepithelial cysts while maintaining mucociliary differentiation (Figure 2B) (1), 5 out of 6 TC+P xenografts showed subclonal squamous differentiation, dyskeratosis, and absence of mucociliary markers (Figure 2A, 2F, Supplementary Figures 2B and 3A, B). TC+PS xenografts displayed a similar phenotype. Previously, we reported subclonal *SOX2* expression in TC+PS ALI cultures due to negative selection of SOX2-overexpressing (*SOX2*^oe^) cells in the context of *PTEN* truncation (1). Thus, the squamous phenotype in TC+PS xenografts could reflect enhanced fitness of *SOX2*^oe^ cells in vivo. To test this, we compared mCherry (co-expressed with *SOX2* via a P2A element; Supplementary Figure 3C) immunostaining intensity between squamous and mucociliary areas in TC+PS xenografts (Figure 2G). No significant differences in mCherry levels were detected (Supplementary Figure 3D), indicating that squamous differentiation is not due to selection against *SOX2*^oe^ cells and likely arises from the same mechanism in both TC+P and TC+PS xenografts (Figure 2G).

In our prior work, we used the apical expansion of p63-positive cells as a marker of LUSC development in ALI cultures (1). Due to architectural differences in xenografts, we assessed the overall frequency of p63-positive cells instead. No significant correlation was found between p63+ cell frequencies in xenografts and ALI cultures. TC+P and TC+PS mutants behaved as outliers (Supplementary Figure 3E), likely due to their aberrant squamous differentiation. Removing these outliers revealed a significant positive correlation among the remaining genotypes (Supplementary Figure 3F).

In conclusion, we observed several differences between mutant hBEC phenotypes in vivo and in vitro. TC+PK xenografts grew larger than expected but retained a mucociliary, cystic architecture. The most notable divergence was the emergence of subclonal squamous differentiation in TC+P and TC+PS xenografts, contrasting with the mucociliary phenotype in ALI cultures. This difference may reflect prolonged in vivo growth and selective pressures. Nonetheless, these variations do not contradict our previous conclusions regarding the cooperative role of the three pathways in driving aggressive LUSC phenotypes and the proposed evolutionary trajectory—initiated by *SOX2*^oe^-driven squamous metaplasia and followed by sequential activation of the Nrf2 and PI3K/Akt pathways.

### Gene expression

To further investigate differences in the behaviour of our LUSC model in vivo and in vitro, we performed bulk RNA sequencing on matched xenografts and ALI cultures. Due to the lack of engraftment of the wild-type and TC mutants, the limited size of the TC+S and TC+KS xenografts, and poor RNA quality in some replicates, we successfully sequenced xenografts for five genotypes: TC+P (n=5), TC+K (n=5), TC+PS (n=4), TC+PK (n=5), and TC+PKS (n=6).

Principal component analysis (PCA) of the transcriptomes revealed that samples clustered primarily by genotype, although a secondary separation based on experimental condition (xenograft vs. ALI culture) was also apparent (Figure 3A), particularly among TC+PKS xenografts. As in our previous study (1), clustering was primarily influenced by the presence of *KEAP1* mutations (bottom clusters) and *SOX2* overexpression (right cluster). Overall, TC+K and TC+PK xenografts and ALI cultures showed close clustering, indicating similar transcriptional profiles between the two systems. In contrast, TC+PKS samples showed a more pronounced divergence between xenograft and ALI conditions. Additionally, some TC+P and TC+PS xenografts exhibited outlier behaviour, shifting rightward on the PCA plot, likely reflecting varying degrees of squamous differentiation (Figure 2A and 2F). Notably, these outlier xenografts clustered closer to TC+S ALI cultures, which exhibit a prominent squamous phenotype (1).

**Figure 3.**
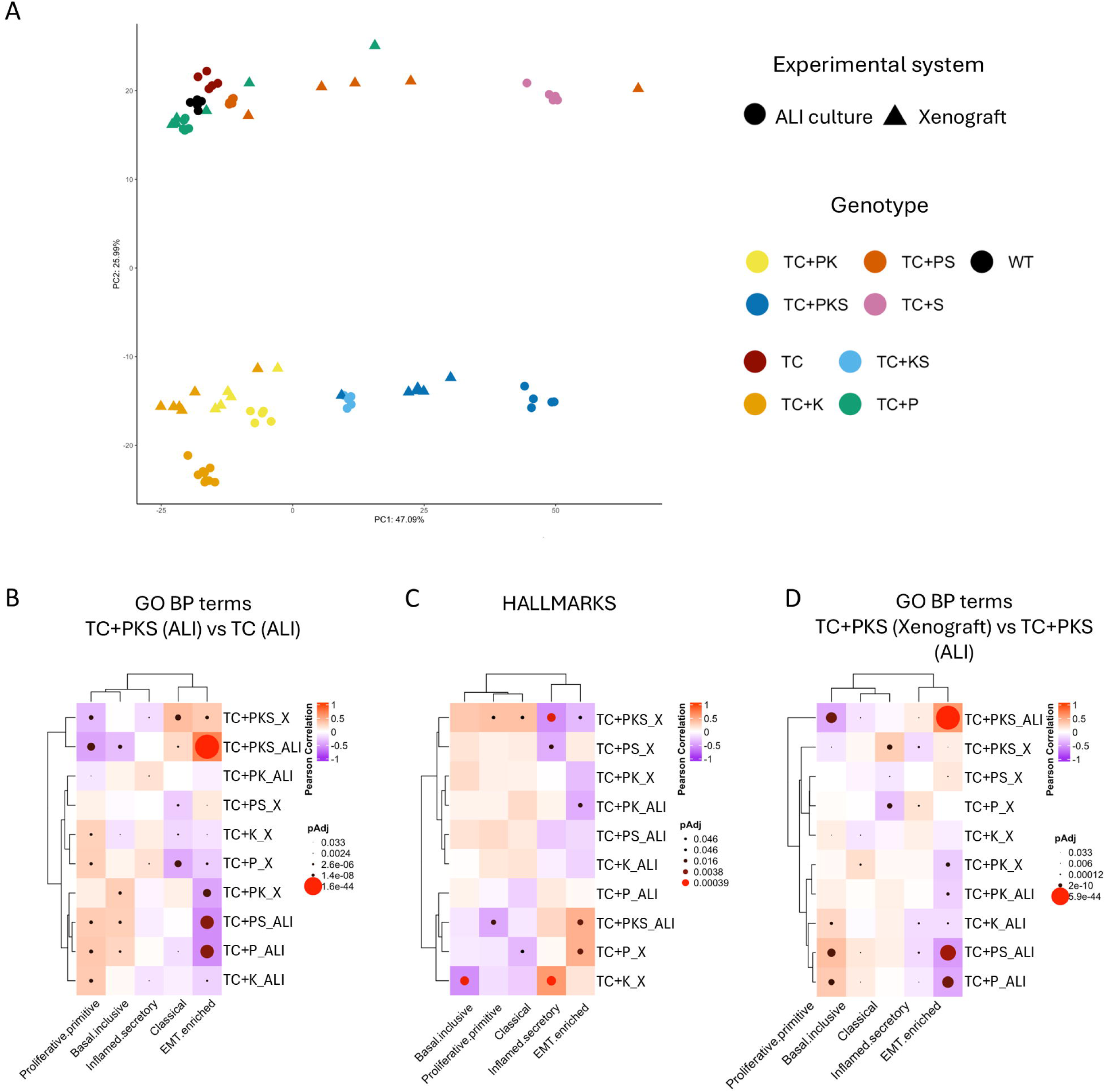
Comparative gene-expression analysis between xenografts and ALI cultures generated with mutant hBECs from the same donor. **A.** Principal component analysis including ALI cultures and xenografts. We also included ALI culture samples with genotypes that did not grow in vivo (wild type and TC) or were too small for transcriptomic analysis (TC+S and TC+KS). **B, C and D.** Correlation analysis between the normalised enrichment scores (NES) of molecular signatures calculated using ssGSEA in LUSC subtypes (horizontal) and xenografts or ALI cultures (vertical). **Panel B** includes GO Biological Processes significantly regulated by the simultaneous activation (TC+PKS vs TC comparison) of the squamous differentiation, Nrf-2 and PI3k/Akt pathways in ALI cultures as reported in Ogden et al., (2025). **Panel C** include the Hallmarks Collection (MSigDB). **Panel D** includes GO Biological Processes significantly enriched in the xenograft vs ALI comparison for the TC+PKS mutant. Correlation coefficients and adjusted p-values were calculated using Pearson analysis in the three panels.

To identify the nature of the transcriptomic differences between experimental systems, we conducted differential expression analysis between xenografts and ALI cultures for the TC+PKS genotype (Supplementary Data 1), followed by gene set enrichment analysis (GSEA; Supplementary Data 2 and 3) and GO Biological Process (GO BP) term condensation using semantic similarity (Tables 1 and 2, Supplementary Data 4 and 5). We focused on the TC+PKS genotype because it exhibited the most pronounced in vivo vs. in vitro transcriptional differences in PCA and represents the most aggressive genotype in both contexts. However, differential expression results for all sequenced genotypes (xenograft vs. ALI) are available in Supplementary Data 6–9.

**Table 1.** Clusters of semantically related GO Biological Processes significantly enriched in upregulated genes (adjusted p-value<0.05) in the TC+PKS xenografts vs TC+PKS ALI cultures. Significantly enriched GO Biological Processes were identified using gene-set enrichment analysis (adjusted p-value<0.05) and clusters of semantically related terms were identified using the simplifyEnrichment package.

| Cluster number | Summary terms | # GOBP terms |
| --- | --- | --- |
| 1 | Sperm cell development | 5 |
| 2 | Neuron/axon/ photoreceptor cell/ dendrite formation | 11 |
| 3 | Stem cell maintenance | 4 |
| 4 | Eye/ear morphogenesis, kidney development | 17 |
| 5 | Patterning | 5 |
| 6 | Cilium assembly | 9 |
| 7 | Cilium/cell projection assembly | 8 |
| 8 | Cell junction assembly | 5 |
| 9 | Chromatin/ chromatid/DNA conformation | 7 |
| 10 | Microtubule formation | 10 |
| 11 | DNA damage/double-strand break repair | 8 |
| 12 | Epigenetics, histone modification | 16 |
| 13 | Sperm cell flagellum assembly | 1 |
| 14 | Sensory/mechanical perception | 2 |
| 15 | Metal detoxification | 6 |
| 16 | Tissue homeostasis | 5 |
| 17 | Membrane repolarisation | 7 |
| 18 | Synaptic transmission | 5 |
| 19 | Cytoskeleton, smoothened pathway, chromosome organisation | 10 |
| 20 | Cilium localisation | 6 |
| 21 | Cilium transport | 2 |
| 22 | Cilium/flagellum motility | 5 |
| 23 | Cytoskeleton related transport | 9 |
| 24 | Cilium/cytoskeleton movement | 6 |

**Table 2.** Clusters of semantically related GO Biological Processes significantly enriched in downregulated genes (adjusted p-value<0.05) in the TC+PKS xenografts vs TC+PKS ALI cultures. Significantly enriched GO Biological Processes were identified using gene-set enrichment analysis (adjusted p-value<0.05) and clusters of semantically related terms were identified using the simplifyEnrichment package.

| Cluster | Summary terms | # GO BP terms |
| --- | --- | --- |
| 1 | Amino acid transport | 3 |
| 2 | Virus-host interaction | 7 |
| 3 | Epithelia/fat/blood cell differentiation | 15 |
| 4 | Development | 18 |
| 5 | Development, stem cell maintenance | 9 |
| 6 | Cholesterol/alcohol metabolism | 16 |
| 7 | Nucleoside metabolism | 11 |
| 8 | Amino acid/lipid metabolism | 9 |
| 9 | Metabolism | 31 |
| 10 | Autophagy | 6 |
| 11 | Protein degradation | 18 |
| 12 | mRNA stability, translation, transcription factor-DNA binding | 13 |
| 13 | Protein modification | 19 |
| 14 | Protein phosphorylation | 25 |
| 15 | Apoptosis, MAPK pathway, JNK signalling | 17 |
| 16 | Apoptosis, Notch pathway, epidermal growth factor pathway, ER stress, | 35 |
| 17 | Response to hormones, response to unfolded protein | 25 |
| 18 | Response to external stimulus/nutrients/osmosis | 6 |
| 19 | Response to external stimulus/nutrients/osmosis/unfolded protein | 17 |
| 20 | Cytokine production, IL-2 production | 8 |
| 21 | Miscellaneous | 13 |
| 22 | Muscle adaptation, response to ROS, chemotaxis | 9 |
| 23 | Intracellular transport, Golgi, ER, lipid transport | 8 |
| 24 | Intracellular transport, regulation of protein localisation | 20 |
| 25 | Lamellipodium assembly | 7 |
| 26 | Cell junction assembly, cell-substrate junction | 14 |
| 27 | Positive regulation of cell death | 6 |
| 28 | Miscellaneous | 13 |
| 29 | Negative regulation of cell growth, cell-cell and cell-matrix adhesion | 13 |

In the TC+PKS xenograft vs. ALI comparison, we identified 4,014 differentially expressed genes (adjusted p-value < 0.05), of which 2,695 were upregulated (logLFC 0.33 to 48) and 1,319 were downregulated (|logLFC| 0.28 to 6.83) (Supplementary Data 1). GSEA followed by GO BP term clustering revealed 24 positively enriched and 29 negatively enriched clusters (Tables 1 and 2, Supplementary Data 4 and 5).

Positively enriched GO BP clusters included terms related to cilium assembly, flagellum function, axon and microtubule organization, and cytoskeletal remodelling (Table 1, Supplementary Data 4), suggesting increased migratory or motility capacity in TC+PKS xenografts. This phenotype may result from physical constraints on cell movement imposed by ALI culture inserts, or from interactions with the mouse stroma in vivo. Alternatively, it may reflect localized mucociliary differentiation. Although some regions of TC+PKS xenografts exhibited apical acetylated-tubulin staining indicative of ciliation (Supplementary Figure 4A), expression levels of canonical markers for ciliated (*FOXJ1*) and goblet cells (*MUC5AC*) did not significantly differ between xenografts and ALI cultures (Supplementary Figures 4B and 4C).

Additional positively enriched GO BP clusters were associated with DNA double-strand break repair, developmental processes, stem cell maintenance, and histone modification (Table 1, Supplementary Data 4). In contrast, negatively enriched clusters encompassed diverse biological processes, including apoptosis, post-translational modification, intracellular transport, protein degradation, and metabolism of amino acids, nucleosides, and lipids (Table 2, Supplementary Data 5).

A key goal of this transcriptomic analysis was to assess whether in vivo transcriptomes more closely resemble patient tumours compared to ALI cultures. To evaluate this, we conducted correlation analyses similar to those from our previous study (1). Specifically, we extracted all GO BP terms significantly regulated by the three oncogenic pathways targeted in our model (TC+PKS ALI vs. TC ALI) and calculated Pearson correlations between the normalized enrichment scores (NES) of these terms in xenografts and ALI cultures, and in human LUSC molecular subtypes (9) using ssGSEA (Figure 3B).

This analysis revealed a significant and positive correlation between the TC+PKS transcriptome and the classical LUSC subtype for both xenografts and ALI cultures. However, this correlation was stronger in xenografts (ALI vs. classical: Pearson r = 0.22, adjusted p = 1.49 × 10⁻L; xenograft vs. classical: Pearson r = 0.39, adjusted p = 1.09 × 10⁻¹¹) (Figure 3B). To confirm this finding using an unbiased set of gene sets, we repeated the analysis using the Hallmarks gene set collection (Figure 3C). This revealed no significant correlation between ALI cultures and the classical subtype (Pearson r = –0.12, adjusted p = 0.56), while the TC+PKS xenografts again showed a significant positive correlation (Pearson r = 0.37, adjusted p = 0.04). Notably, TC+PKS xenografts also correlated significantly with the proliferative-primitive LUSC subtype (Pearson r = 0.36, adjusted p = 0.04) (Figure 3C).

Finally, we tested the hypothesis that NES values for GO BP terms significantly enriched in the xenograft vs. ALI comparison for TC+PKS correlate with the classical LUSC subtype. This would support the idea that observed in vivo–in vitro differences are biologically meaningful rather than technical artifacts, and that xenografts more accurately model LUSC biology. Consistent with this, we observed a non-significant correlation between TC+PKS ALI cultures and the classical subtype (Pearson r = 0, adjusted p = 0.37), but a significant and positive correlation in TC+PKS xenografts (Pearson r = 0.30, adjusted p = 1.97 × 10^−10^;) (Figure 3B).

In summary, our transcriptomic analysis confirmed that the presence of *SOX2* overexpression and *KEAP1* truncation was the primary driver of transcriptomic clustering—consistent with our in vitro findings (1). However, we also identified transcriptional differences attributable to the experimental context, particularly in the TC+PKS genotype. These differences, notably the upregulation of genes involved in motility and development, suggest that the gene expression of xenografts better recapitulate the biology of classical LUSC tumours.

### Identification of new drivers of squamous differentiation in LUSC

To uncover mechanisms driving squamous differentiation in the TC+P and TC+PS genotypes— independent of *SOX2*^oe^—we hypothesized that this phenotype could arise from somatic alterations positively selected in vivo. To test this, we performed whole-exome sequencing (WES) on xenografts and their matched pre-injection cells, aiming to identify mutations present exclusively in xenografts. As with RNA-seq, sample size limitations restricted WES analysis of xenografts to five genotypes: TC+P (n=6), TC+K (n=5), TC+PS (n=5), TC+PK (n=6), and TC+PKS (n=6). Pre-injection cells were sequenced for all genotypes.

To identify candidate drivers of squamous differentiation, we first generated a list of somatic mutations present in xenografts but absent from their corresponding pre-injection cells, excluding silent and intronic variants (Supplementary Data 10). We then focused on mutations found in both TC+P and TC+PS xenografts but absent in other genotypes (Supplementary Figure 5A, Supplementary Data 11). This filtering strategy yielded 14 genes mutated in at least three TC+P or TC+PS xenografts (Figure 4A). Notably, the same mutation was present in each of the 14 genes across all relevant samples (Figure 4A, Supplementary Data 11), suggesting that these alterations were pre-existing at low frequency in the TC+P population prior to the generation of TC+PS mutants via lentiviral transduction.

**Figure 4.**
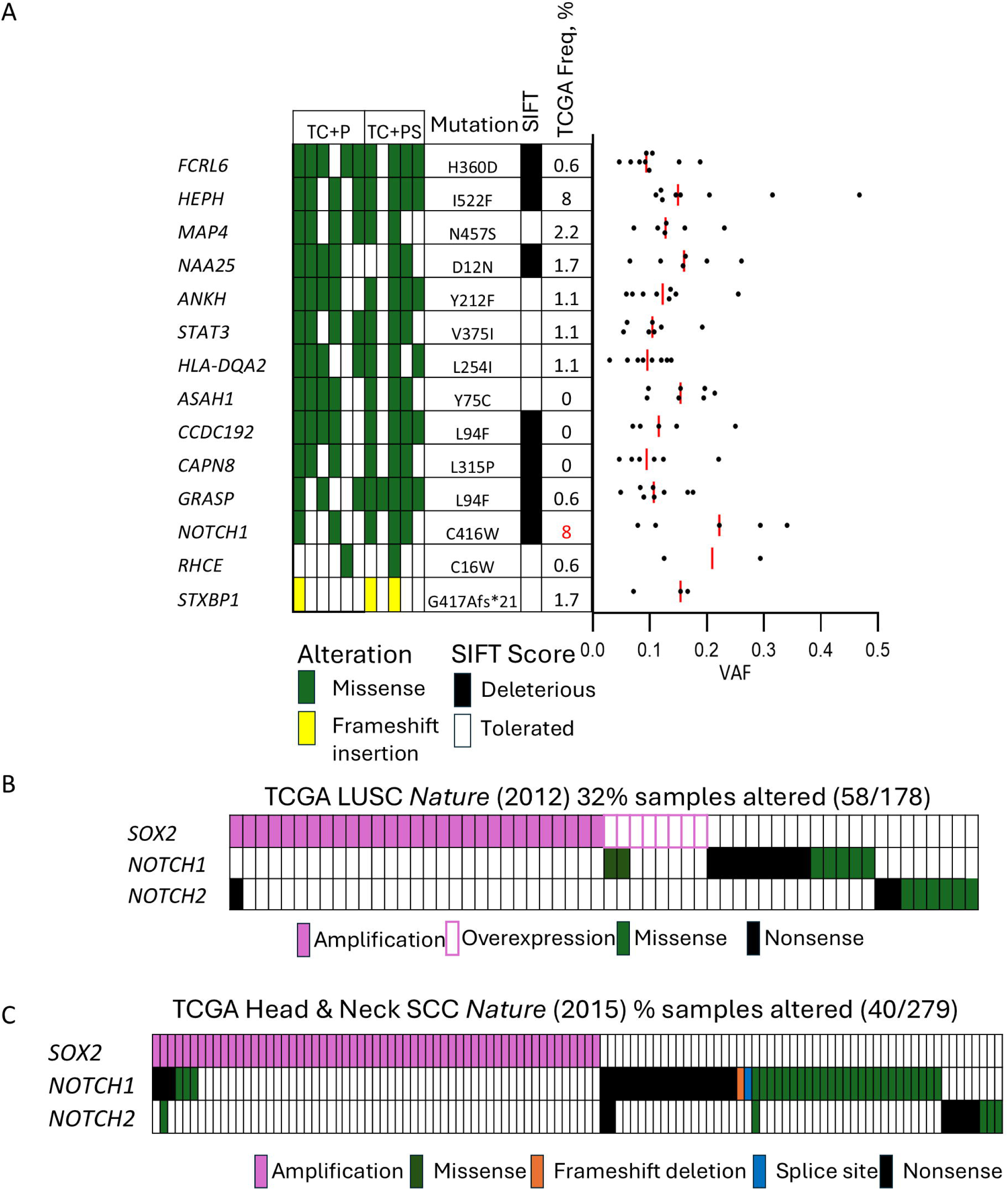
Whole-exome sequencing analysis of xenografts with spontaneous squamous differentiation (TC+P and TC+PS). **A.** Panel showing non-synonymous mutations present in TC+P and TC+PS xenografts and absent in the genotype-matched pre-injection hBECs and parental wild-type hBECs. The panel shows the amino acid change, SIFT scores, percentage of TCGA samples with mutations in the same genes (red indicate that the gene is significantly mutated in the TCGA cohort) and variant allelic frequencies (VAFs) in xenografts (red lines shows median VAFs). **B and C**. *SOX2*, *NOTCH1* and *NOTCH2* somatic alterations in the LUSC and head and neck squamous carcinomas cohorts as published by the TGCA consortium in Nature (2012 and 2015 respectively). Only tumours with alteration in at least one of the three genes are shown.

Among these, a *NOTCH1* missense mutation (C416W) emerged as the strongest candidate driver of the squamous phenotype. This variant was present in 2 of 6 TC+P and 3 of 5 TC+PS xenografts (Figure 4A) and was predicted to be deleterious to protein function. *NOTCH1* was the only significantly mutated gene among the 14 candidates based on the TCGA LUSC cohort (8), and the C416W variant had the highest median variant allele frequencies (VAFs) (Figure 4A). Manual inspection of sequencing reads confirmed that this mutation was also present in 3 additional TC+P and 1 TC+PS xenografts at subclonal levels (VAF <10%) (Supplementary Figure 5B).

Importantly, the TCGA data indicate that *NOTCH1* and *NOTCH2* mutations show minimal overlap with *SOX2* amplification or overexpression in LUSC (Figure 4B) [5], suggesting mutually exclusive roles. A similar pattern is observed in head and neck squamous cell carcinoma (HNSCC), another cancer type where mutations in the NOTCH pathway have been implicated in tumour initiation (10, 11), and where *NOTCH1/2* mutations also rarely coincide with *SOX2* amplification (Figure 4B) (12).

Although the specific C416W mutation has not been reported in the LUSC or HNSCC TCGA cohorts, the majority of *NOTCH1* missense mutations in these cancers are unique. This mutational heterogeneity is consistent with the tumour suppressor role of *NOTCH1* being compromised by a broad spectrum of missense variants.

Taken together, these observations implicate *NOTCH1* C416W as a potential driver of squamous differentiation in TC+P and TC+PS xenografts, suggesting that loss of *NOTCH1* function promotes squamous transdifferentiation in LUSC independently of *SOX2* amplification.

### *NOTCH1* truncation induces squamous differentiation and impairs mucociliary identity in ALI cultures

Based on the above findings, we hypothesized that *NOTCH1* loss-of-function might directly induce squamous differentiation in the absence of SOX2 overexpression. To test this, we generated human basal epithelial cells (hBECs) with CRISPR/Cas9-mediated truncation of *NOTCH1* in the context of *TP53* and *CDKN2A* loss (TC+N), using the same rationale described in our previous work (1). Specifically, exon 8 of *NOTCH1* was targeted to introduce frameshift indels near codon 422 (Supplementary Figure 6A). We then compared the phenotypes of wild-type, TC, TC+N, and TC+S ALI cultures.

Histological examination of ALI cultures revealed that TC+N mutants exhibited areas with disrupted pseudostratified morphology and loss of mucociliary differentiation, displaying instead squamous characteristics with flattened, multilayered cells (Supplementary Figure 6B, Figure 5A). However, the squamous phenotype was more pronounced in the TC+S mutant, which showed thicker epithelial layers and extensive shedding of squamous cells (Figure 5A).

**Figure 5.**
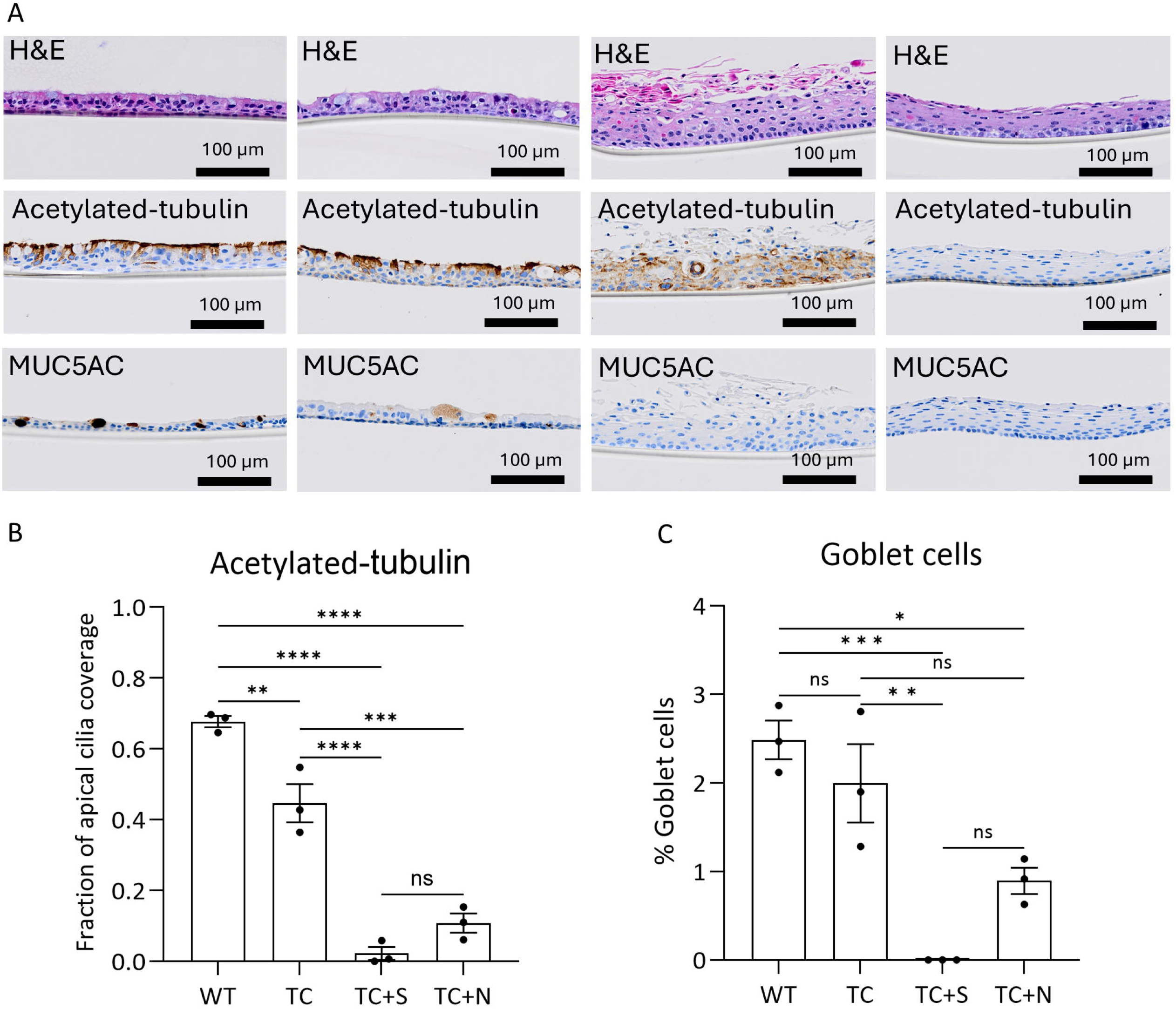
Comparative phenotypic analysis of *NOTCH1* truncation in ALI cultures. **A.** Representative images of histological sections from wild type, TC, TC+S and TC+N ALI cultures stained with H&E (top row), MUC5AC (middle row) and acetylated-tubulin (bottom row). N=*NOTCH1* truncation **B and C.** Quantification of ciliated (B) and goblet cells (C) in wild type, TC, TC+S and TC+N ALI cultures. Ciliated cells were quantified as the fraction of apical acetylated-tubulin, and goblet cells as the percentage of MUC5AC-positive cells. Data shown as mean±SEM. Adjusted p-values were calculated by one-way ANOVA, followed by Kruskal-Wallis test for multiple comparison correction. Statistical significance shown as: ns= not significant ∗p < 0.05, ∗∗p < 0.01, ∗∗∗p < 0.001, and ∗∗∗∗p < 0.0001

To quantify these observations, we assessed mucociliary differentiation using immunohistochemical markers: acetylated-tubulin for ciliated cells and MUC5AC for goblet cells (Figure 5A, B). TC+N mutants displayed a marked reduction in ciliated cells compared to wild-type and TC mutants, although not to the extent observed in TC+S. Goblet cell frequency was also reduced in TC+N mutants relative to wild-type, but comparable to TC cultures.

Overall, these results confirm that *NOTCH1* loss-of-function impairs mucociliary differentiation and promotes squamous features, even in the absence of *SOX2* overexpression. Although the phenotype is less severe than in TC+S mutants, these findings support a model in which *NOTCH1* inactivation constitutes an alternative driver of the squamous differentiation pathway in LUSC.

## DISCUSSION

This study was designed with two primary objectives: a) to determine whether our mutant hBEC-based LUSC models yield consistent results in vivo and in vitro, and b) to highlight the specific strengths of each approach.

Our previous work using ALI cultures demonstrated that the sequential activation of squamous differentiation, Nrf-2, and PI3K/Akt pathways drives progression from normal bronchial epithelium to premalignant squamous stages, culminating in aggressive LUSC phenotypes (1). This evolutionary trajectory aligned closely with the classical LUSC subtype. Our in vivo results mirrored these findings.

While wild-type and TC mutants failed to grow in vivo, TC+S xenografts formed small, stratified, polarized nodules exhibiting extensive dyskeratosis. TC+KS xenografts were similarly small but showed more disorganized architecture and expansion of p63-positive cells. In contrast, TC+PKS xenografts retained a disorganized structure but were significantly larger than TC+KS nodules and more invasive. Although TC+PK and TC+PKS xenografts achieved comparable volumes, TC+PK nodules were cystic and maintained a highly organized and benign mucociliary morphology.

Gene expression analyses showed that the mutant hBEC transcriptomes were more strongly influenced by *SOX2*^oe^ and *KEAP1* truncation than by the experimental platform (ALI culture vs xenograft). This dependency had already been observed in our previous in vitro analyses (1). Nonetheless, the experimental system also contributed to transcriptional variability, especially among TC+PKS mutants, largely through genes associated with microtubule formation and cell motility. These differences improved alignment with the classical LUSC subtype transcriptome.

In addressing our first objective, we conclude that phenotypic outcomes in vivo and in vitro are broadly concordant with respect to evolutionary trajectories. However, xenograft implantation produces tumours that more closely mimic the classical LUSC subtype at the gene expression level.

These transcriptomic differences may reflect the physical constraints on cell migration inherent to ALI cultures, cues from the in vivo microenvironment, or both. A hybrid system allowing for migration while preserving organotypic structure could help clarify the role of cell motility. Likewise, co-culture with stromal components may better model in vivo conditions.

Our in vivo analysis identified a tumour-suppressive role for *NOTCH1* in LUSC, based on the phenotypic divergence between TC+P and TC+PS mutants. Loss of *NOTCH1* function promoted a squamous phenotype in the absence of *SOX2* amplification. The presence of a *NOTCH1* mutation (C416W) in TC+P and TC+PS xenografts, specifically in the context of *PTEN* truncation, raises the possibility of cooperation between NOTCH pathway inhibition and PI3K/Akt activation in driving a subset of *SOX2*-independent LUSC. However, this potential synergy remains hypothetical and requires functional validation. The co-occurrence may also be incidental, as the C416W mutation could have arisen in other genotypes by chance.

Moreover, the low individual frequencies of *NOTCH1* and *PTEN* mutations in LUSC (8), despite being significantly mutated genes, limit the ability to analyse co-mutation patterns at the population level. Still, prior functional studies have independently supported a tumour-suppressive role for *NOTCH1* in LUSC (13), and *NOTCH1* mutations have been associated with downstream signalling changes in this cancer (9). However, these studies did not explore the relationship between *NOTCH1* mutations and the absence of *SOX2* amplification.

Regardless this newly described role for *NOTCH1* mutations in LUSC initiation, our findings underscore a key strength of the xenograft system: its capacity to uncover novel LUSC drivers under selective pressures in vivo, complementing the more controlled but limited in vitro system. Expanding the genetic diversity of mutant hBECs may further enhance this advantage by fuelling Darwinian selection. Such diversity could be introduced by modelling known sources of genomic instability in LUSC, including genome-doubling, APOBEC activity, and cigarette smoke exposure (14). Alternatively, targeting epigenetic regulators frequently altered in LUSC, such as *ARID1A* and *KMT2D*, may induce transcriptional diversity, providing a substrate for selective adaptation (8, 9, 15).

Despite these strengths, the in vitro organotypic system remains highly valuable due to its versatility, lower cost, ability to assess cell-intrinsic effects of candidate LUSC drivers and the identification of downstream processes that we validated in patient tumours (1). Moreover, it enables the analysis of genotypes that do not form tumours in xenografts.

In conclusion, we do not advocate exclusively for either organotypic cultures or xenografts as the superior system for modelling LUSC progression. Rather, the choice of model should depend on the specific research question and available resources. Our data support the use of genetically engineered hBECs in vitro as a robust and flexible platform to elucidate mechanisms of LUSC progression using a hypothesis-driven approach to validate candidate drivers and in vivo as this approach enables unbiased discovery.

## MATERIALS AND METHODS

### Cell culture

Mouse embryonic 3T3-J2 fibroblasts (Kerafast, EF3003) were maintained in DMEM (Gibco™, 11995065) supplemented with 10% bovine serum (Fisher Scientific, 16-107-078), 2 mM GlutaMAX™ (Gibco™, 35050061), and 5% Penicillin-Streptomycin (Gibco™, 15070063). Lenti-X 293T cells (Takara, 632180) and primary human lung fibroblasts (Lonza Bioscience, ID: HLF-2, batch: 20TL356516) were cultured in DMEM (Gibco™, 11995065) containing 10% fetal bovine serum (LabTech, 80837), 2 mM GlutaMAX™, and 5% Penicillin-Streptomycin.

Normal human bronchial epithelial cells (hBECs) were purchased from Lonza (CC-2540S). For the side-by-side comparative study in xenografts and ALI cultures we used batch# 18TL290281 (75-year-old, black male, non-smoker). For the phenotypic analysis of *NOTCH1* truncation in ALI cultures we used batch# 482214 (62-years-old, caucasian male, non-smoker)

hBECs were cultured as previously described (16). Briefly, sub-confluent 3T3-J2 fibroblasts were treated with 4 µg/ml Mitomycin C (Sigma-Aldrich, M4287) for 2 hours, washed three times with PBS, trypsinized (Gibco™, 12605036), and seeded at 20,000 cells/cm². The culture medium was then replaced with a 1:3 mixture of Ham’s F-12 Nutrient Mix (Gibco™, 11765054) and DMEM (Gibco™, 41966), supplemented with 10% fetal bovine serum (Gibco™, 11563397), 5 µM Y-27632 dihydrochloride (Bio-Techne Sales Corp, 1254), 25 ng/ml hydrocortisone (Sigma-Aldrich, H0888), 0.125 ng/ml EGF (Thermo Fisher Scientific, 10-605-HNAE50), 5 μg/ml insulin (Sigma-Aldrich, I6634), and 0.1 nM cholera toxin (Sigma-Aldrich, C8052). Human bronchial epithelial cells were seeded at 20,000 cells/cm². Fresh 3T3-J2 fibroblasts were used for each epithelial cell passage. All cultures were maintained in a humidified incubator at 37°C with 5% COL. Media was changed every two days, and all cell lines were routinely screened for mycoplasma contamination.

### Genome editing and plasmid construction

To design sgRNAs for this study, we utilized the Benchling platform (Benchling.com), which provides sgRNA candidates along with on-target and off-target scoring. For each gene, we selected sets of four sgRNAs with the lowest off-target scores, prioritizing those that targeted regions within or upstream of the first annotated functional domain in the open reading frame (ORF) to ensure effective protein inactivation. To minimize the risk of unintended editing at other loci, we adopted an approach similar to that described by Drost et al. (4), sequencing the top five predicted off-target sites with the highest off-target scores using Sanger sequencing. The CRISPR-Cas9 target sites selected are listed in the table below:

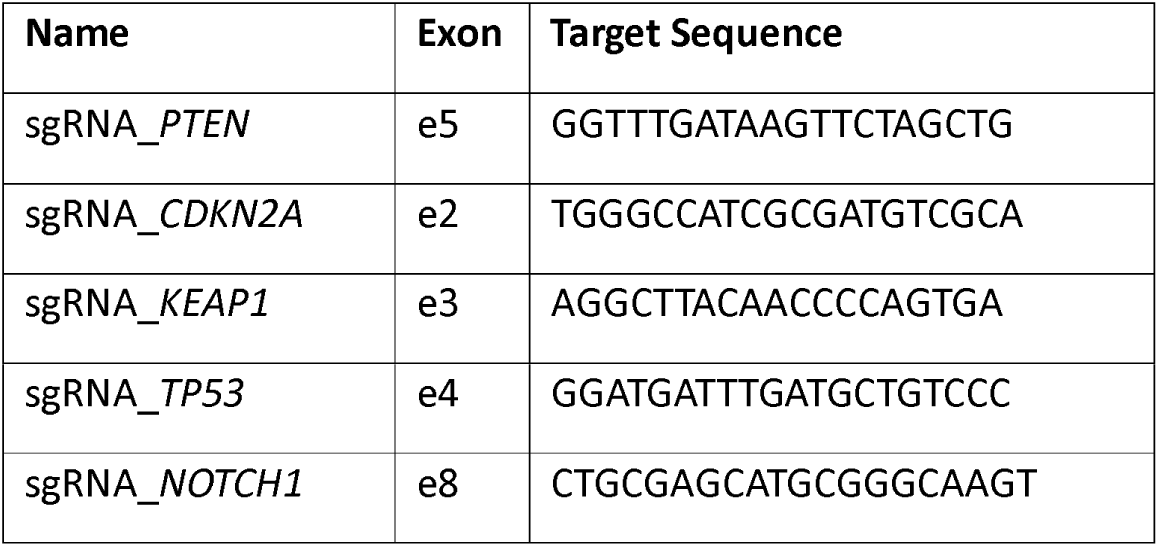

Multiplex CRISPR-Cas9 plasmids targeting *TP53*, *CDKN2A*, *PTEN*, *KEAP1*, and *NOTCH1* were generated using the one-step CRISPR-Cas9 assembly kit (Addgene Kit #1000000055), kindly provided by Takashi Yamamoto (17). Guide RNAs were incorporated into pX330 plasmid backbones through Golden Gate cloning, following established protocols (17).

To induce *SOX2* overexpression, we employed the third-generation lentiviral vector pUltra-hot (Addgene #24130), generously shared by Malcolm Moore. SOX2 coding sequence was PCR-amplified with flanking restriction sites—EcoRI at the 5′ end (GAATTC) and BbsI at the 3′ end (GAAGAC)—and cloned into the multiple cloning site of the pUltra-hot vector via restriction digestion and ligation, resulting in the construct pUltra-hot–SOX2.

### Bronchial epithelial cell electroporation and selection

Multiplex CRISPR-Cas9 pX330 plasmids were introduced into normal bronchial epithelial cells via electroporation using the P3 Primary Cell 4D-Nucleofector® X Kit L (Lonza, V4XP-3024) in conjunction with the 4D-Nucleofector™ X Unit (Lonza). A total of 500,000 cells were pelleted and resuspended in 100 µl of Nucleofector™ Solution, followed by the addition of 5 µg of pX330 plasmid DNA. The entire mixture was transferred to a Nucleocuvette™ vessel and subjected to electroporation using the DC-100 program. After electroporation, cells were incubated at room temperature for 10 minutes, then diluted with 500 µl of pre-warmed medium before being seeded into T25 flasks containing 4.5 ml of culture medium and a feeder layer of mitotically inactivated mouse embryonic 3T3 fibroblasts prepared as previously described (11).

Cells were allowed to recover for 48 hours before initiating selection for TP53 mutants by adding Nutlin-3A (Sigma-Aldrich, SML0580) to a final concentration of 5 nM. Selection was maintained for five days, with media replacement and fresh Nutlin-3A added every two days. Typically, colonies of mutant bronchial epithelial cells became visible 2 to 3 days after completing the selection process.

### Bronchial epithelial cell transduction

Stable overexpression of *SOX2* in bronchial epithelial cells was achieved via lentiviral transduction using viral particles carrying the packaged pUltra-hot–SOX2 construct. Lentivirus production was performed by transfecting Lenti-X 293T cells at 90% confluency with 880 ng of pUltra-hot–SOX2, 572 ng pMDLg/pRRE (Addgene #122251), 308 ng pCMV-VSV-G (Addgene #8454), and 220 ng pRSV-Rev (Addgene #12253), all in 2 ml of culture medium with 6 µl of FuGENE HD Transfection Reagent (Promega, #E2311). Viral particles were concentrated using a polyethylene glycol (PEG) solution (40% w/v) containing 2.4% NaCl in sterile water. Viral titers (copies/ml) were quantified with the Lenti-X™ qRT-PCR Titration Kit (Takara Bioscience, 631235), following the manufacturer’s protocol.

An identical procedure was used to produce control lentivirus particles containing the empty pUltra-hot vector (lacking *SOX2* cDNA), which served as a negative control. For transduction, 250,000 bronchial epithelial cells were seeded into T25 flasks containing 5 ml of medium and a feeder layer of mitotically inactivated 3T3-J2 mouse fibroblasts, prepared as described previously (16). While the epithelial cells were still in suspension, a transduction mixture—comprising 4 × 10L viral copies, 12 µg/ml polybrene, and PBS to a final volume of 100 µl—was added to the culture. This protocol consistently achieved transduction efficiencies exceeding 90%. After 48 hours, the medium was replaced, and transduction efficiency was confirmed by flow cytometry via detection of mCherry fluorescence.

### PCR genotyping

For genotyping to identify CRISPR-Cas9-induced indels, genomic DNA was extracted from mutant bronchial epithelial cells using proteinase K digestion. Approximately 500,000 cells were incubated in 500 µl of proteinase K digestion buffer (50 mM KCl, 50 mM Tris-HCl, 2.5 mM EDTA, 0.45% NP40, and 0.45% Tween) along with 50 µg of Proteinase K (ThermoFisher Scientific, EO0491) at 55°C for 5 hours. The enzyme was then inactivated by heating the samples at 95°C. Primers for PCR amplification were as follows: CDKN2A_forward 5’-CACCCTGGCTCTGACCATTC-3’, CDKN2A_reverse 5’-GCAAGTCCATTTCGGGATTA-3’, KEAP1_forward 5’-TACGACTGCGAACAGCGAC-3’, KEAP1_reverse 5’-GGCACAGAATCAAAGGTCAC-3’, PTEN_forward 5’-TCCAGTGTTTCTTTTAAATACCTGTT-3’, PTEN_reverse 5’-GGGGGAGAATAATAATTATGTGAGGT-3’, TP53_forward 5’-CAGGAAGCCAAAGGGTGAA-3’, TP53_reverse 5’-CCCATCTACAGTCCCCCTTG-3’, NOTCH1_forward 5’-GAGTGCTCGCTGGGTAGG and NOTCH1_forward 5’-AAGCAACCCACAGATGTTCC.

### In vivo tumourigenesis

All animal procedures were approved by the Animal Welfare and Ethics Review Body at Cancer Research UK Manchester Institute and carried out by members of the Biological Resources Unit Core Facility. To preserve blinding, the operators were not informed of the experimental design, the rationale of the experiment or the genotypes of the injected cells. A total of 2×10^6^ hBECs of the same passage (p9) were resuspended in 1:1 RPMI: Cultrex (Trevigen, 3432-010-01) and injected subcutaneously into the right flank of 8-to 12-week-old female NSG mice (Envigo) (n=6 mice per genotype, 9 genotypes). Animals were identified with numerical IDs unrelated with genotypes and allocated to cages that contained animals of two different genotypes for randomisation purposes. Mice were housed under pathogen-free conditions, 12 hour light-dark cycle, in individually ventilated cages maintained at 20-24°C and 40-60% relative humidity and fed with standard diet. Xenograft size was measured using a digital calliper. Mice were removed from the study 105 days following injection, except for one TC+PK replicate that was culled on day 89 due to reaching the maximum size. We did not exclude any mice from the study. Visible tumours were harvested and divided into two halves: one for fixation in formalin for FFPE tissue and another for snap freezing for later by DNA (DNeasy Blood & Tissue Kit, Qiagen, 69504) and RNA extraction (RNeasy Mini Kit, Qiagen, 74104). Small xenografts that could not be divided were fixed in formalin for FFPE tissues only.

### Air-liquid interface culture and processing

Air–liquid interface (ALI) organotypic cultures were established using wild type and mutant human bronchial epithelial cells. Cells were grown on Falcon® Permeable Supports with 0.4 µm pore-size PET membranes (Corning, 353095). To prepare the membranes, each insert was coated with 20 µg of type I bovine collagen. Specifically, a 3 mg/ml PureCol® solution (CellSystems, 5005) was diluted 1:30 in sterile PBS, and 200 µl was added to each membrane, followed by a 1-hour incubation at room temperature. After incubation, the collagen solution was removed, and the membranes were gently rinsed with PBS.

A total of 30,000 bronchial epithelial cells were pelleted and resuspended in 100 µl of Airway Epithelial Cell Growth Medium (PromoCell, C-21160), then seeded directly onto the collagen-coated membranes, which were placed in individual wells of a 24-well plate. The basal compartment was filled with 500 µl of the same medium. Cells were expanded under submerged conditions for 7 days, with medium changes every 2 days.

On day 7, the apical medium was removed to initiate air–liquid interface conditions, and the basal compartment was replenished with 500 µl of PneumoCult™-ALI Maintenance Medium (STEMCELL Technologies, 05100). Cultures were maintained for 21 days to allow for epithelial differentiation, with medium in the basal compartment replaced every 2 days. Upon completion, cultures were harvested for downstream analyses.

For histological evaluation, cultures were fixed in 4% paraformaldehyde for 30 minutes, embedded in agarose, and processed into paraffin blocks for sectioning. For gene expression analysis, cultures were rinsed with PBS and total RNA was isolated using the RNeasy Mini Kit (Qiagen, 74104) according to the manufacturer’s protocol.

### Immunofluorescence and immunohistochemistry

4µm FFPE sections were cut from FFPE blocks from ALI organotypic cultures and xenografts. Immunohistochemistry was used for anti-TTF1 (Abcam, ab76013) (1:100), anti-p63 (Abcam, ab124762) (1:400) antibodies, anti-human mitochondria (Abcam, ab92824) (1:500), anti-acetylated tubulin (Sigma, T6793) (1:3000), anti-MUC5AC (Invitrogen, MA5-12178) (1:4500) and anti-mCherry (Novus, NBP2-25157) (1:500). Antigen retrieval (ER 1, 20 minutes) and staining was performed using the Leica Bond RX automated platform using the classical IHC-F protocol with antigen retrieval achieved using BOND™ Epitope Retrieval solution 1 (Lecia, AR9961) for 20 minutes. Acetylated tubulin and MUC5AC staining protocols included an additional 30-minute blocking step with casein (Sigma-Aldrich, B6429). Chromogenic 3,3-diaminobenzidine (DAB) staining was achieved using the BOND™ Polymer Refine Detection Kit (Lecia, DS9800).

### Image acquisition and analysis

All histological slides were scanned at 20x magnification using the Olympus VS200 brightfield scanner with built-in Köhler and LED illumination and an integrated 2/3 inch CMOS camera. Image analysis was performed with the QPath software (version 0.5.1), utilizing the DAB module to upload images. The percentage of MUC5AC- and p63-positive cells was determined by calculating the ratio of positively stained cells to the total number of nuclei within each defined analysis region using the default settings of *Positive cell detection* tool. For acetylated tubulin, coverage was assessed by measuring manually the length of epithelium stained positively and dividing it by the total epithelial length evaluated. mCherry expression was quantified as mean DAB staining intensity for a pixel size=2 μm using the *Add intensity features* tool.

### RNA-sequencing

Total RNA was extracted using the RNeasy Mini Kit (Qiagen, 74104) following the manufacturer’s protocol. RNA library preparation and sequencing were performed by the Molecular Biology Core Facility at the Cancer Research UK Manchester Institute. Xenograft samples were homogenised prior to extraction. Only samples with RNA Integrity Numbers (RIN) greater than 7 were selected for sequencing. Libraries were prepared from 100 ng of total RNA using the Lexogen QuantSeq 3′ mRNA-Seq Library Prep Kit.

Sequencing was carried out on an Illumina NovaSeq 6000 platform using 100 bp single-end reads. Base calling was followed by conversion to FASTQ files via Illumina’s bcl2fastq software. Adapter sequences were trimmed using the autodetect function in Trim Galore (v0.6.5), and reads were aligned to the GRCh38 reference genome using the STAR aligner (v2.5.1b). Resulting BAM files were quantified in R (v3.6.1) using the featureCounts function from the Rsubread package (v2.0.1).

Principal component analysis (PCA) was performed using regularised log-transformed read counts generated by DESeq2 (v1.26). Surrogate variable analysis (SVA) was applied to detect and adjust for hidden batch effects or confounding variables in the combined dataset. Differential expression analysis was conducted with DESeq2, incorporating SVA-derived covariates into the model to ensure robust comparisons between experimental conditions.

### Gene-set correlation analysis and correlation analyses between LUSC samples and mutant hBECs

Pre-ranked Gene Set Enrichment Analysis (GSEA) was performed for pairwise comparisons using the fgsea package (v1.24.0), based on gene-level statistics from DESeq2. Genes were filtered for an adjusted p-value (pAdj)<0.05 and ranked by logL fold change. To reduce redundancy among Gene Ontology (GO) terms, we applied the simplifyEnrichment tool (18).

To evaluate transcriptomic similarity between lung squamous cell carcinoma (LUSC) molecular subtypes and mutant human bronchial epithelial cells (hBECs) both in vivo (xenografts) and in vitro (ALI cultures), we obtained data from the CPTAC-LUSC cohort (9) via the GDAC portal. Transcript per million (TPM, unstranded) gene expression data were assembled into a unified count matrix, and patient metadata were retrieved from the supplementary materials of the original study. Samples labelled as “mixed subtype” were excluded, and mean expression values were computed for each gene within individual molecular subtypes.

Single-sample GSEA (ssGSEA) was used to calculate normalised enrichment scores (NES) for three curated sets of gene signatures:

1. GO Biological Process terms significantly enriched in the TC+PKS (ALI) vs TC (ALI) comparison, as defined in Ogden et al. (2025) (1);
2. The Hallmark gene sets from MSigDB (https://www.gsea-msigdb.org/gsea/msigdb/human/genesets.jsp?collection=H);
3. GO Biological Process terms significantly enriched in TC+PKS (xenograft) vs TC+PKS (ALI) comparisons.

For the first and third gene set collections, enrichment terms were derived from pre-ranked GSEA using DESeq2 output from the corresponding comparisons.

These gene sets were then used in ssGSEA analyses, applied to both the averaged gene expression profiles from the CPTAC-LUSC cohort and the mean profiles from our ALI and xenograft samples. Correlations between NES values from CPTAC-LUSC subtypes and hBEC-derived ALI cultures were quantified using Pearson’s correlation. P-values were adjusted for multiple testing using the Benjamini-Hochberg procedure.

### Whole exome sequencing

Whole exome sequencing data were acquired using the Twist whole exome V2 kit using standard methodology and sequenced using Illumina Novoseq 6000 (paired-end, 100 bp read length, 200x coverage). Basecalls were produced using bcl2fastq (version 2.20.0.422). Raw reads were trimmed using trim galore (version 0.6.10) and mapped to both human (B37; GATK) and mouse (mm10; Ensembl release 102). Human data were separated from possible mouse contamination using XenofilteR (version 1.6). Extracted human alignments were subject to cleaning and recalibration as prescribed by Broad using GATK (version 4.0.4). Somatic SNV calls were produced using Mutect2. Calls were filtered using FilterMutectCalls and subset to Twist probe locations. Filtered somatic calls were converted to MAF format using vcf2maf (version 1.6.21) and analysed using MAFtools (version 2.16.0).

### Quantification and Statistics

Statistical testing of tumour size and immunostaining was carried out using the GraphPad Prism 10.4.1 software. Statistical comparisons were performed using a one-way ANOVA with *post hoc* tests for multiple comparison correction. Pairwise comparisons were carried out using Student’s t-test for normally distributed data or Mann-Whitney test for non-normally distributed data. Correlation analyses were carried out using Pearson analysis. Significance was determined by p value < 0.05. Figures denote significance where ∗p < 0.05, ∗∗p < 0.01, ∗∗∗p < 0.001, and ∗∗∗∗p < 0.0001. Data was presented as means with standard deviation. n denotes the number of replicates.

### Data availability

Accession number for xenograft DNA-seq and RNA-seq: GSE302966

Accession number for ALI RNA-seq: PRJNA1043668 (Donor 1, Lonza ID: 36722)

## AUTHOR CONTRIBUTIONS

JO: Conceptualization, data curation, formal analysis, investigation, methodology, validation, visualization and writing-original draft, review & editing

RS: Conceptualization, data curation, formal analysis, methodology, software, validation

SS: Conceptualization, data curation, formal analysis, methodology, software, supervision and validation

AO: Data curation, investigation, methodology, project administration, resources, validation and writing-review & editing

CD: Conceptualization, funding acquisition, methodology, resources, supervision and writing-review & editing

CLG: Conceptualization, data curation, formal analysis, funding acquisition, methodology, project administration, resources, supervision, validation, visualization and writing-original draft, review & editing

## ACKNOWLEDGEMENTS

This project has been funded by the National Centre for the Replacement, Refinement & Reduction of Animals in Research (NC/W001284/1), the Rosetrees Trust (M767), the Cancer Research UK Lung Cancer Centre of Excellence (A25146) and the Cancer Research UK Manchester Institute (C5759/A20971).

The animal experiments were carried out under Home Office project licence PP4143150.

Genetic manipulations were carried out under approval by Genetic Modification and Biohazards Safety Committee (University of Manchester).

The authors used generative artificial intelligence (ChatGPT) for language correction.

We would like to thank the Computational Biology Support, Molecular Biology, Histopathology, Biological Resources Unit and Scientific Computing core facilities (grant no. C5759/A27412) at the Cancer Research UK Manchester Institute and the National Biomarker Centre.

## SUPPLEMENTARY FIGURE LEGENDS

**Supplementary Figure 1:**
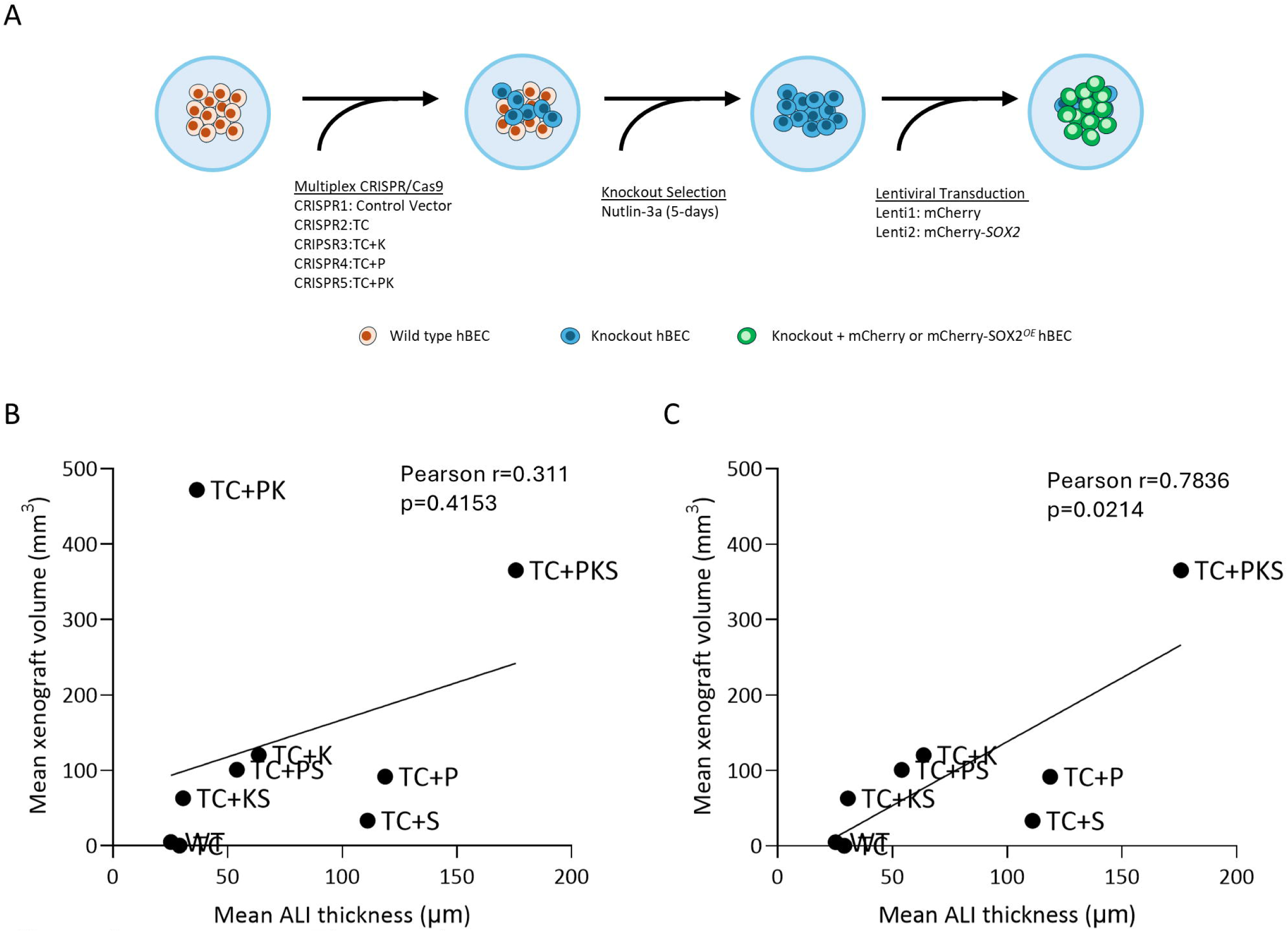
Protocol for genetic manipulation of hBECs and correlation between xenograft volume and ALI thickness. **A.** Schematics depicting the protocol of hBEC genetic manipulation using CRISPR/Cas9 and lentiviral transduction. Wild type hBECs were electroporated to transfect multiplex CRISPR constructs and treated with Nutlin-3a to select for *TP53* truncations as all constructs include a *TP53*-targeting gRNA. Nutlin-3a resistant hBECs were transduced with polycistronic lentiviral constructs that harbour mCherry+*SOX2* or mCherry only. **B.** Correlation between mean xenograft volume and mean ALI thickness for each genotype. Line shows a linear regression of the data. Correlation coefficient, and the p-value was calculated using Pearson analysis. **C.** Correlation between mean xenografts volume and mean ALI thickness for each genotype except TC+PK. Line shows a linear regression of the data. Correlation coefficient and p-value were calculated using Pearson analysis.

**Supplementary Figure 2.**
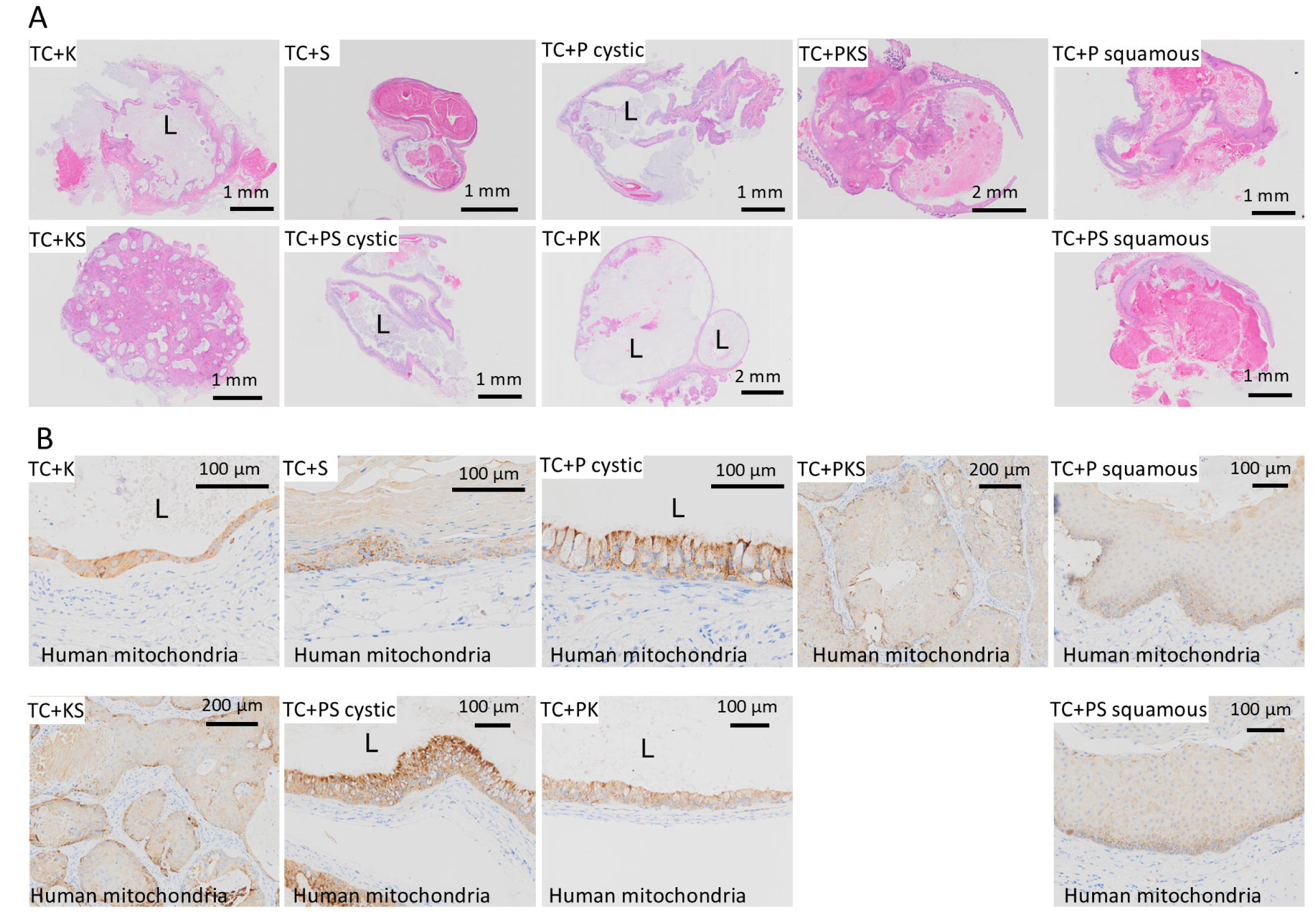
Histological analysis of xenografts generated with mutant hBECs. **A.** Representative H&E images of the histological structure of xenografts at low magnification. For TC+P and TC+PS mutants, representative images of xenografts with predominant squamous and cystic morphologies are shown separately. L=Lumen. **B.** Representative images of human mitochondria immunostaining of xenografts to identify the presence of human cells. For TC+P and TC+PS mutants, representative images of areas with squamous and cystic morphologies are shown separately. L=Lumen.

**Supplementary Figure 3.**
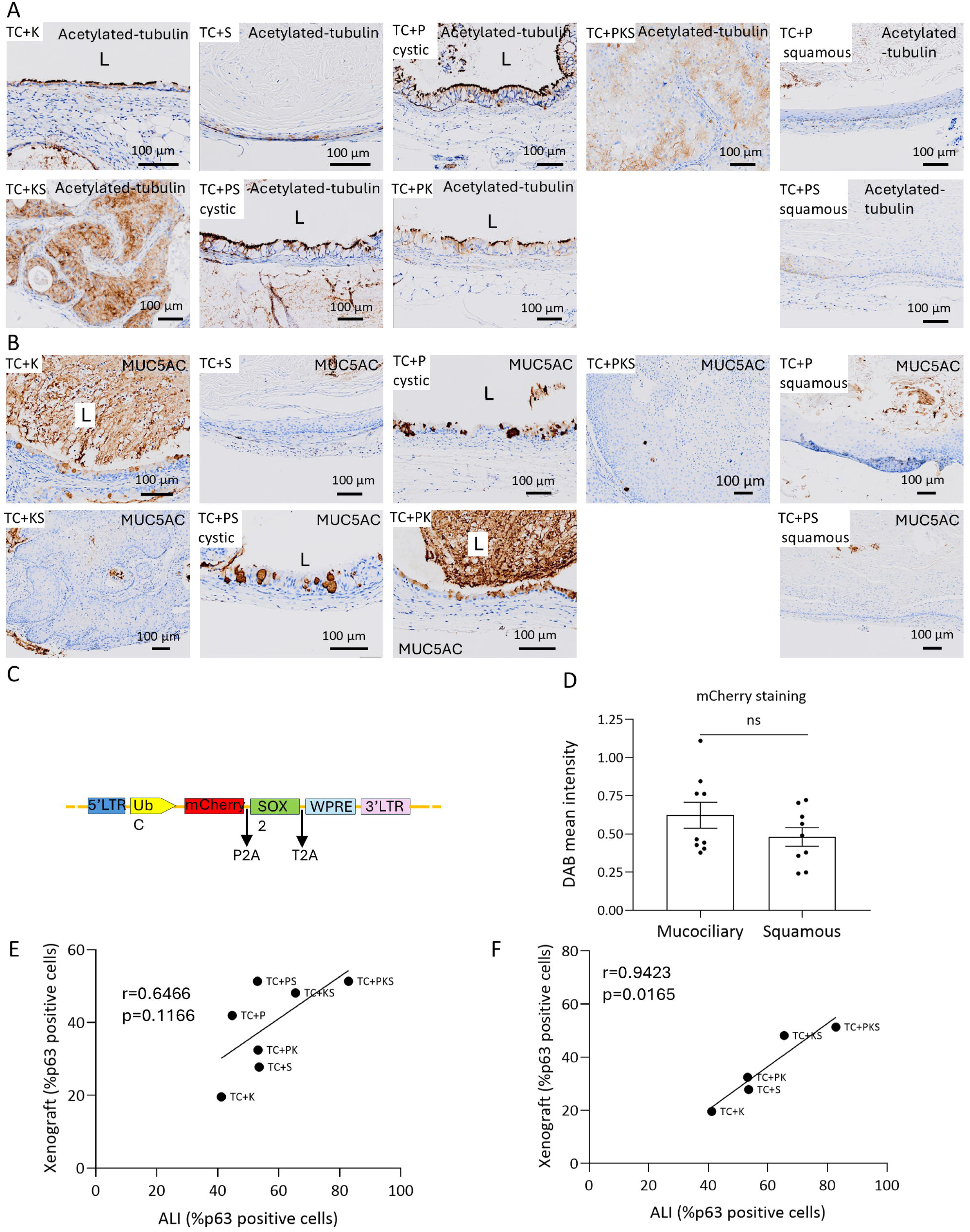
Histological analysis of xenografts generated with mutant hBECs (continued). **A and B:** Representative images of acetylated-tubulin and MUC5AC immunostaining of xenografts to identify the presence of ciliated (**A**) and goblet cells (**B**) respectively. For TC+P and TC+PS mutants, representative images of areas with squamous and cystic morphologies are shown separately. L=Lumen. **C.** Schematic depicting the expression cassette of the lentiviral construct pUltrahot for *SOX2* overexpression. 5’LTR and 3’LTR= 5’ and 3’ long terminal repeats; UbC=Ubiquinone promoter; WPRE= Woodchuck Hepatitis Virus Posttranscriptional Regulatory Element; P2A and T2A=ribosome skipping sequences. **D.** Quantification of mCherry staining in areas juxtaposed mucociliary and squamous morphology in TC+PS xenografts (see Figure 2F). mCherry staining was quantified as brown DAB mean pixel intensity for each region analysed. Data is show as mean±SEM (n=9). P-values were calculated using a paired t-test. **E and F.** Correlation analysis between the frequency of p63-positive cells in xenografts and ALI cultures for each genotype. Panel E includes all genotypes that generated xenografts, whereas panel F excludes the outlier TC+P and TC+PS genotypes. Line shows a linear regression of the data. Correlation coefficient and the p-value were calculated using Pearson analysis. ns=not significant

**Supplementary Figure 4.**
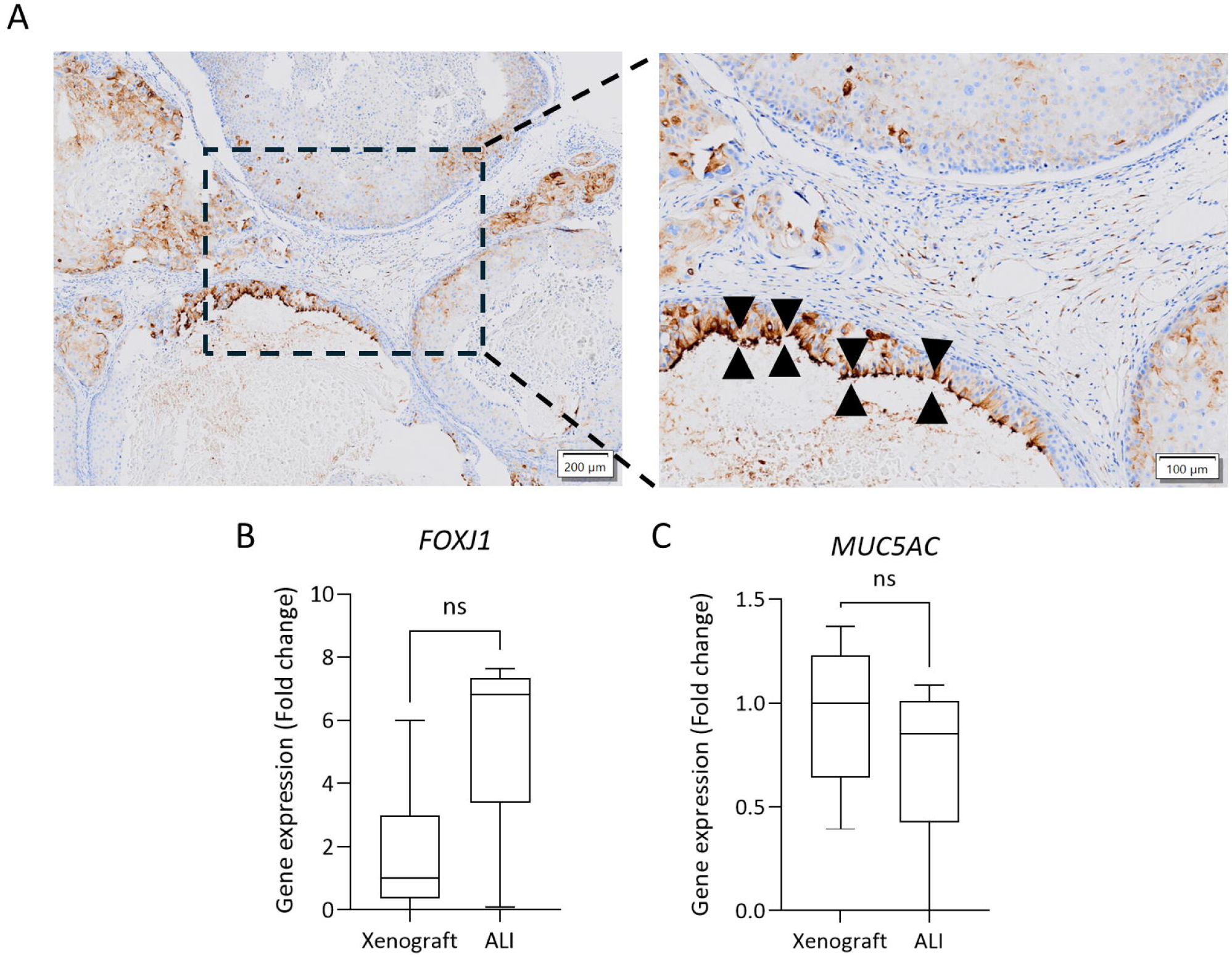
Analysis of mucociliary differentiation in TC+PKS xenografts. **A.** Acetylated-tubulin-immunostaining to identify the presence of ciliated cells in TC+PKS xenografts. **B and C**. Comparative analysis of the expression of transcriptomic markers of ciliated (*FOXJ1*) **(B)** and goblet cells (*MUC5AC*) GO Biological Processes **(C)** in ALI cultures and xenografts using RNA-seq data (Fold-change of normalised TPM). Data shown as median, interquartile range (Q1-Q3) and 5-95 percentile (whiskers). p-values were calculated using Mann-Whitney test. ns=not significant

**Supplementary Figure 5.**
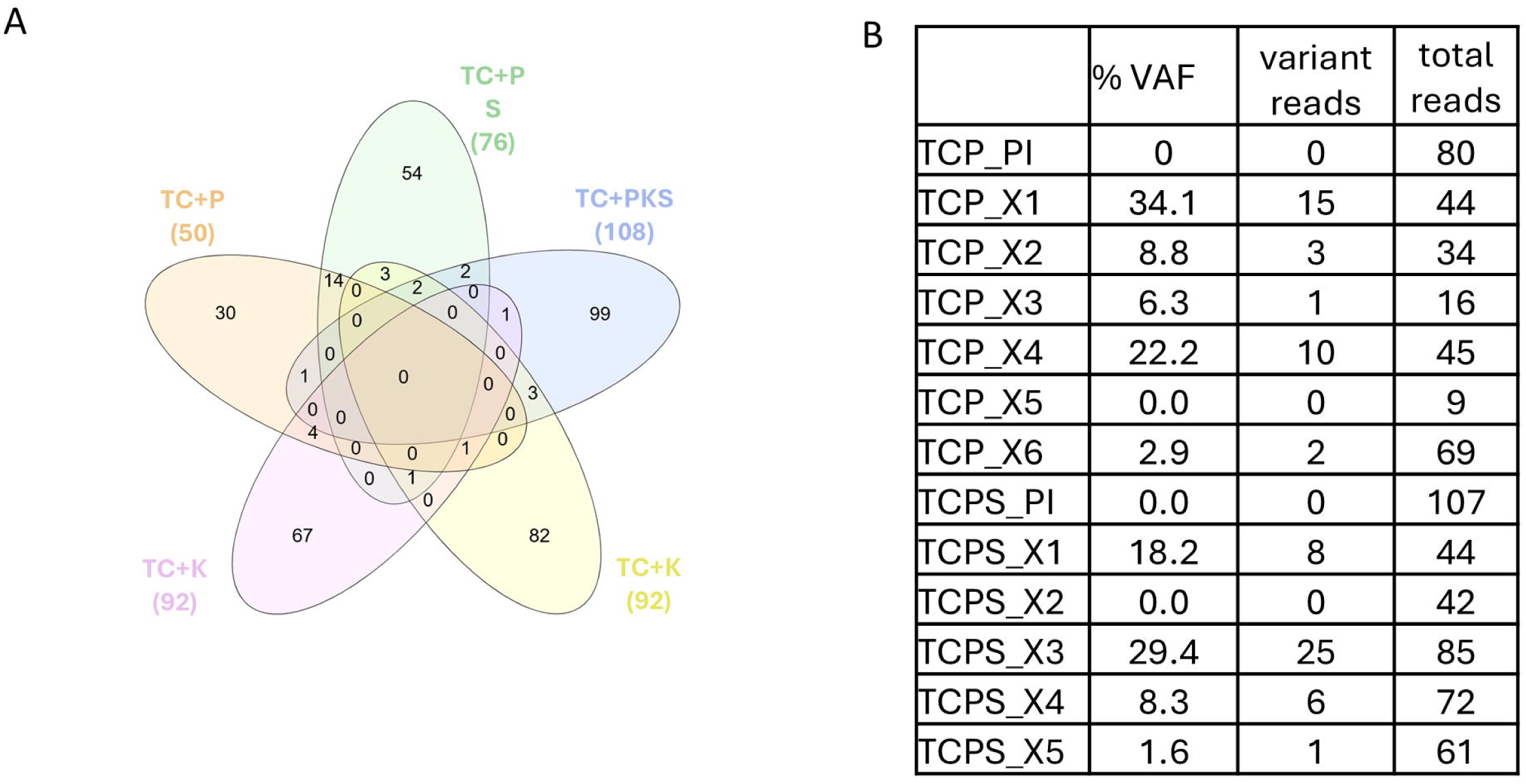
Whole-exome sequencing analysis of xenografts with spontaneous squamous differentiation (TC+P and TC+PS). Continued from Figure 4. **A.** Venn diagram depicting the overlap between genes mutated in xenografts of each genotype analysed by whole-exome sequencing. Mutations present in the pre-injection population were excluded. **B**. Table showing the variant allelic frequencies (VAFs) for the *NOTCH1* C416W mutation in TC+P and TC+PS xenografts. PI=pre-injection population; X=xenograft.

**Supplementary Figure 6.**
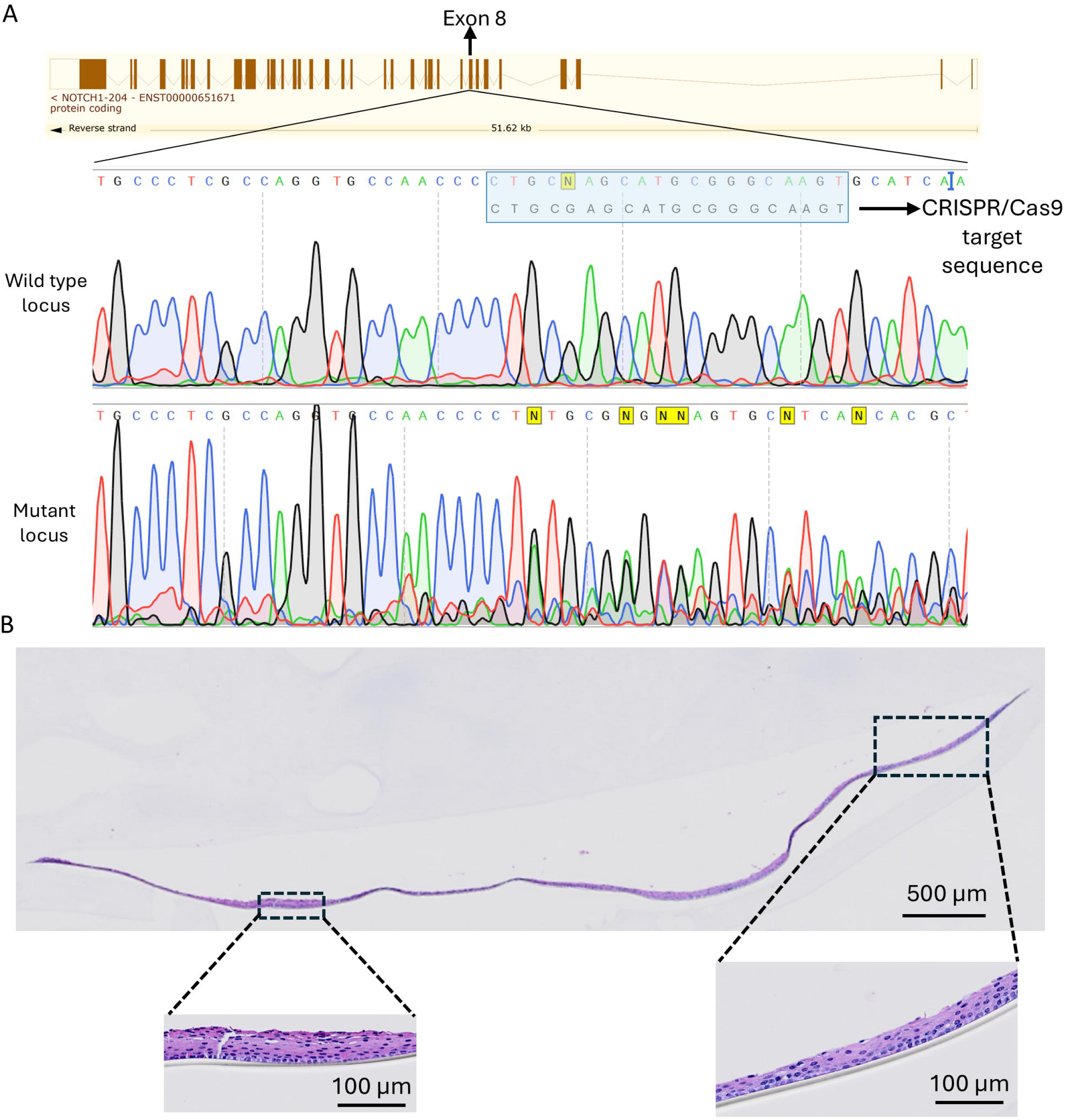
Phenotypic analysis of *NOTCH1* truncation in ALI cultures. **A.** Sanger sequencing of the CRISPR/Cas9 targeted *NOTCH1* locus in cells transfected with empty vector and cells transfected with the *NOTCH1* targeting construct. The gRNA targeted codon 422 in exon 8. **B.** Low magnification image of an H&E-stained TC+N ALI cultures highlighting some areas with loss of pseudostratified morphology.

## SUPPLEMENTARY DATA LEGENDS

**Supplementary Data 1.** Differentially expressed genes in the TC+PKS xenograft vs. TC+PKS ALI culture comparison (adjusted p-value<0.05).

**Supplementary Data 2.** GO Biological Process, GO Molecular Function and GO Cell Compartment gene-sets significantly enriched (q<0.05) in the list of upregulated genes (adjusted p-value<0.05) in the TC+PKS xenograft vs. TC+PKS ALI culture comparison.

**Supplementary Data 3.** GO Biological Process, GO Molecular Function and GO Cell Compartment gene-sets significantly enriched (q<0.05) in the list of downregulated genes (adjusted p-value<0.05) in the TC+PKS xenograft vs. TC+PKS ALI culture comparison.

**Supplementary Data 4**. Semantically related clusters of GO Biological Process in the list of upregulated genes (adjusted p-value<0.05) in the TC+PKS xenograft vs. TC+PKS ALI culture comparison.

**Supplementary Data 5**. Semantically related clusters of GO Biological Process in the list of downregulated genes (adjusted p-value<0.05) in the TC+PKS xenograft vs. TC+PKS ALI culture comparison.

**Supplementary Data 6, 7, 8 and 9.** Differentially expressed genes (adjusted p-value<0.05) in the xenograft vs ALI culture comparison for the TC+K, TC+P, TC+PK and TC+PS genotypes.

**Supplementary Data 10.** List of somatic mutations in xenografts and matched pre-injection mutant hBECs. Somatic mutations were selected based of absence of mutant reads in the wild-type parental population. Each tab shows one genotype.

## REFERENCES

1. Ogden J, Sellers R, Sahoo S, Oojageer A, Chaturvedi A, Dive C, et al. A human model to deconvolve genotype-phenotype causations in lung squamous cell carcinoma. Nat Commun. 2025;16(1):3215.

2. Matano M, Date S, Shimokawa M, Takano A, Fujii M, Ohta Y, et al. Modeling colorectal cancer using CRISPR-Cas9-mediated engineering of human intestinal organoids. Nat Med. 2015;21(3):256–62.

3. Hodis E, Torlai Triglia E, Kwon JYH, Biancalani T, Zakka LR, Parkar S, et al. Stepwise-edited, human melanoma models reveal mutations’ effect on tumor and microenvironment. Science. 2022;376(6592):eabi8175.

4. Drost J, van Jaarsveld RH, Ponsioen B, Zimberlin C, van Boxtel R, Buijs A, et al. Sequential cancer mutations in cultured human intestinal stem cells. Nature. 2015;521(7550):43–7.

5. Daniel VC, Marchionni L, Hierman JS, Rhodes JT, Devereux WL, Rudin CM, et al. A primary xenograft model of small-cell lung cancer reveals irreversible changes in gene expression imposed by culture in vitro. Cancer Res. 2009;69(8):3364–73.

6. Guillen KP, Fujita M, Butterfield AJ, Scherer SD, Bailey MH, Chu Z, et al. A human breast cancer-derived xenograft and organoid platform for drug discovery and precision oncology. Nat Cancer. 2022;3(2):232–50.

7. Xu X, Kumari R, Zhou J, Chen J, Mao B, Wang J, et al. A living biobank of matched pairs of patient-derived xenografts and organoids for cancer pharmacology. PLoS One. 2023;18(1):e0279821.

8. Cancer Genome Atlas Research N. Comprehensive genomic characterization of squamous cell lung cancers. Nature. 2012;489(7417):519–25.

9. Satpathy S, Krug K, Jean Beltran PM, Savage SR, Petralia F, Kumar-Sinha C, et al. A proteogenomic portrait of lung squamous cell carcinoma. Cell. 2021;184(16):4348–71 e40.

10. Loganathan SK, Schleicher K, Malik A, Quevedo R, Langille E, Teng K, et al. Rare driver mutations in head and neck squamous cell carcinomas converge on NOTCH signaling. Science. 2020;367(6483):1264–9.

11. Nyman PE, Buehler D, Lambert PF. Loss of Function of Canonical Notch Signaling Drives Head and Neck Carcinogenesis. Clin Cancer Res. 2018;24(24):6308–18.

12. Cancer Genome Atlas N. Comprehensive genomic characterization of head and neck squamous cell carcinomas. Nature. 2015;517(7536):576–82.

13. Sinicropi-Yao SL, Amann JM, Lopez DLY, Cerciello F, Coombes KR, Carbone DP. Co-Expression Analysis Reveals Mechanisms Underlying the Varied Roles of NOTCH1 in NSCLC. J Thorac Oncol. 2019;14(2):223–36.

14. Jamal-Hanjani M, Wilson GA, McGranahan N, Birkbak NJ, Watkins TBK, Veeriah S, et al. Tracking the Evolution of Non-Small-Cell Lung Cancer. N Engl J Med. 2017;376(22):2109–21.

15. Pan Y, Han H, Hu H, Wang H, Song Y, Hao Y, et al. KMT2D deficiency drives lung squamous cell carcinoma and hypersensitivity to RTK-RAS inhibition. Cancer Cell. 2023;41(1):88–105 e8.

16. Hynds RE, Butler CR, Janes SM, Giangreco A. Expansion of Human Airway Basal Stem Cells and Their Differentiation as 3D Tracheospheres. Methods Mol Biol. 2019;1576:43–53.

17. Sakuma T, Nishikawa A, Kume S, Chayama K, Yamamoto T. Multiplex genome engineering in human cells using all-in-one CRISPR/Cas9 vector system. Sci Rep. 2014;4:5400.

18. Gu Z, Hubschmann D. simplifyEnrichment: A Bioconductor Package for Clustering and Visualizing Functional Enrichment Results. Genomics Proteomics Bioinformatics. 2023;21(1):190–202.

